# Slow Oscillations Gate Interictal Spikes Across the Human Thalamocortical–Epileptogenic Network

**DOI:** 10.64898/2026.05.30.728961

**Authors:** Mahmoud Alipour, Wim van Drongelen, Douglas R. Nordli, Joel Voss, Kenneth Lee, David Satzer

**Affiliations:** Department of Neurological Surgery, University of Chicago; Department of Pediatrics, The University of Chicago; Department of Neurology, The University of Chicago

**Keywords:** Slow oscillations, Interictal epileptiform spikes, Focal epilepsy, Closed-loop neuromodulation, Phase–amplitude coupling

## Abstract

**Background:** Slow oscillations (SOs; 0.5–1.5 Hz), a hallmark of non-rapid eye movement (NREM) sleep, are associated with a marked amplification of interictal epileptiform spike (IIS) activity in focal epilepsy. However, the network-level organization of this effect across the thalamocortical–epileptogenic system, and whether IIS-permissive SOs can be predicted from pre-onset brain states, remain unclear.

**Methods:** We analyzed simultaneous scalp EEG and stereo-EEG (SEEG) recordings from 6 patients with drug-resistant focal epilepsy across 24 full-day recording days, sampling prefrontal cortex (PFC), thalamus, and seizure onset zone (SOZ). SO–IIS coupling was characterized across vigilance states using peri-event and phase-based analyses, with a gamma-based validation step to reduce contamination by IIS-related slow potentials. Pre-onset phase-amplitude coupling (PAC) was compared between IIS-permissive and non-permissive SOs.

**Results:** SO–IIS coupling was observed across all regions, with the strongest and most temporally precise pre-trough IIS clustering in the SOZ (peak 4.4% in NREM), exceeding PFC (1.7%) and thalamic coupling. Thalamic coupling was preserved across wakefulness and NREM and was significant in 7/11 nuclei, with nucleus-specific phase preferences. SO morphological features, particularly up-slope and peak-to-peak amplitude at PFC contacts, predicted IIS occurrence in the SOZ. Pre-onset PAC differed significantly between permissive and non-permissive SOs across regions.

**Conclusions:** SO–IIS coupling is a distributed, network-level phenomenon with region- and state-specific characteristics, and pre-onset PAC provides a predictive signature of IIS-permissive brain states. These findings support the feasibility of developing personalized, closed-loop neuromodulatory strategies targeting SO-gated IIS suppression in focal epilepsy.

## 1. Introduction

The link between sleep and epilepsy has been recognized for over a century, yet the exploration of the mechanistic basis of this relationship remains incomplete^1,2^. One of the most consistent clinical observations is the amplification of interictal epileptiform spikes (IIS) during non-rapid eye movement (NREM) sleep, particularly during deep sleep (stage N3), whereas REM sleep is relatively suppressive^3,4^. This sleep-dependent surge in epileptiform activity carries direct clinical consequences across multiple timescales, particularly on cognition. At the short-term level, transient IIS disrupt ongoing cognitive processing, producing reaction time delays even when patients are subjectively unaware of the events^5,6^. Over the course of a night, IIS can interfere with physiological sleep oscillations, including spindles and slow-wave activity, thereby disrupting sleep-dependent memory consolidation and synaptic homeostasis^7,8^. At longer timescales, recurrent interictal activity contributes to network-level dysfunction associated with impaired learning and cognitive performance. Chronically elevated IIS burden is associated with impaired learning, cognitive dysfunction, and developmental consequences, particularly in pediatric epilepsy^9–11^. It is worth noting that the cognitive impact of IIS is not uniform across epilepsy syndromes and may depend on factors beyond IIS burden alone^12^. Beyond cognition, IIS also carry broader clinical relevance as markers of transient epileptogenic network vulnerability and may relate to seizure-related brain states^13^. Collectively, these observations establish sleep-dependent IIS amplification as a clinically significant feature of epilepsy with implications for cognition and epileptogenic network vulnerability. However, despite its clear clinical importance and implications for neuromodulation, the mechanisms by which IIS are temporally organized within brain rhythms—and whether this organization reflects local or distributed network processes, varies across vigilance states, or can be predicted from pre-event neural dynamics—remain largely unresolved.

An important hallmark rhythm of NREM sleep is the slow oscillation (SO; ∼0.5–1.5 Hz), a large-amplitude alternation between up states—periods of sustained neuronal depolarization and firing—and down states—periods of neuronal quiescence and hyperpolarization^14^. SOs are generated primarily within prefrontal cortex (PFC) and coordinated through thalamocortical circuits, propagating as traveling waves across the neocortex^15,16^ and organizing the timing of thalamocortical spindle bursts and hippocampal sharp-wave ripples into a hierarchical temporal structure that supports sleep-dependent memory consolidation^17,18^. The thalamus plays a dual role in this architecture: it contributes to the coordination of cortical SO activity and serves as a major generator of sleep spindles that are nested within SO up states^19,20^. Early work by Steriade and colleagues demonstrated that the synchronized properties of NREM sleep—and slow-wave activity in particular—create conditions that facilitate epileptic discharge propagation^21^. The intuitive mechanism is that the SO up state, characterized by synchronous depolarization of large cortical and subcortical neuronal populations, provides an excitatory context in which pathologically hyperexcitable epileptic networks are more likely to generate IIS^22^.

Direct evidence of SO–IIS temporal coupling at the event level emerged initially from animal and human intracranial studies demonstrating phase-dependent interactions between slow oscillatory activity and epileptiform discharges, where hippocampal and neocortical slow oscillations were shown to facilitate epileptiform events in a state- and phase-dependent manner^23^. In humans, intracranial EEG studies provided the first systematic event-level characterization of SO–IIS coupling, demonstrating that IIS in patients with focal epilepsy preferentially cluster along specific phases of cortical slow oscillations, particularly on the descending limb and near the trough^22^. This phase-dependent coupling has since been replicated and extended across mesial temporal, neocortical, and subcortical regions using both scalp EEG and SEEG recordings^24,25^. Together, these findings establish the SO not merely as a passive background rhythm during which IIS occur, but as an active gating mechanism that coordinates the timing of IIS within the epileptic network.

Despite this progress, the existing literature has predominantly characterized SO–IIS coupling within individual recording sites or limited anatomical regions, leaving the network-level organization of this coupling across the thalamocortical–epileptogenic system poorly understood. The epileptic brain is not a single structure: IIS are generated within and propagated through distributed networks, often involving the seizure onset zone (SOZ), connected cortical regions, and subcortical nodes such as the thalamus. This network framing is supported by recent work showing that IISs can co-occur across spatially separated cortical regions, and that slow oscillations provide a synchronous background that modulates IIS generation and propagation^25^. The thalamus is especially relevant because of its extensive connectivity with cortical networks involved in sleep oscillations^26,27^. More broadly, interictal activity occurs within spatially distributed brain networks that require a system-level approach^10^. IIS can be examined at the level of the SOZ and thalamus using invasive recordings such as stereo-EEG (SEEG), whereas PFC activity can be assessed non-invasively using scalp EEG, particularly from frontal and prefrontal channels. Whether SO–IIS coupling differs in strength, temporal precision, and phase preference across these anatomically and functionally distinct nodes remains unknown, yet clinically relevant: resolving this question would determine whether the SO gating signal is primarily local or network-wide, and which recording modality provides the most informative access to it for future invasive or non-invasive neuromodulation strategies. Furthermore, the nucleus-level organization of thalamic coupling—and whether specific thalamic nuclei with distinct connectivity profiles show differential IIS phase preferences—has not been systematically examined in humans.

A further limitation of the existing literature is its near-exclusive focus on NREM sleep, despite emerging evidence that slow oscillation–like activity during wakefulness can also modulate IIS timing. Intracranial studies have shown that slow oscillation–like waveforms observed during wakefulness share key features with sleep slow oscillations and are associated with fluctuations in network excitability surrounding IIS^28^. These findings suggest that the relationship between slow oscillatory activity and IIS is not restricted to sleep alone, challenging a purely sleep-centric view of SO–IIS coupling. This raises the possibility that IIS gating may reflect a more fundamental property of slow oscillatory dynamics, rather than a mechanism specific to NREM sleep. A direct comparison of coupling strength and temporal organization across wakefulness and NREM sleep within the same thalamic and SOZ recording sites, while controlling for patient and day-level variability, has not been systematically performed.

A significant methodological challenge in SO–IIS coupling studies is the risk of inflating coupling estimates by including IIS-related slow potentials as candidate SOs. IIS can be followed by slow-wave deflections^29^, and morphology-based SO detectors may therefore misclassify some peri-IIS low-frequency events as SOs. Prior studies have addressed this issue primarily through temporal exclusion strategies, for example by removing slow waves occurring in proximity to interictal discharges^22,28^. However, these approaches may obscure the relationship between SOs and IIS by excluding events that are temporally linked, and while they reduce temporal overlap, they do not directly assess whether detected events preserve the physiological properties of canonical slow oscillations. Importantly, experimental and human intracranial studies show that physiological slow oscillations are accompanied by state-dependent modulation of neuronal and high-frequency activity, with reduced gamma activity during the down-state and increased gamma activity during the up-state^22,28,30–33^. Building on this physiological principle, we incorporated a gamma-power validation criterion as part of our analysis of SO detections near IIS events, retaining candidate SOs only when gamma power increased from the trough/down-state to the subsequent peak/up-state.

Beyond characterizing when and where IIS couple to SOs, a key question is whether IIS occurrence can be predicted before the SO unfolds. SOs vary in their association with IIS, with only a subset of events temporally linked to interictal discharges, while others occur without associated epileptiform activity^22^. Here, we refer to these as permissive and non-permissive SOs, respectively. This distinction is particularly important given that SOs are a major target of neuromodulatory interventions aimed at enhancing memory and cognitive function during sleep^34,35^. However, in the epileptic brain, non-selective enhancement of SOs may inadvertently facilitate IIS, introducing a potential trade-off between cognitive benefit and pathological activation. Identifying and predicting SOs that facilitate IIS, as opposed to those that do not and may instead support physiological processes, may therefore be critical for developing safe, selective, and effective closed-loop stimulation strategies that both suppress epileptiform activity and preserve or enhance beneficial slow oscillations.

In the present study, we investigate these questions using simultaneous scalp EEG and SEEG recordings from patients with drug-resistant focal epilepsy, providing access to the thalamocortical–epileptogenic network at three levels: PFC, thalamus, and SOZ. First, we characterize SO–IIS temporal coupling across these regions, examining differences in coupling strength, temporal precision, and phase preference across vigilance states. Second, we assess whether SO morphological features predict IIS occurrence. Third, we analyze the pre-onset network state preceding permissive and non-permissive SOs using phase-amplitude coupling (PAC), mapping the spatial distribution and feature specificity of predictive signals across PFC, thalamus, and SOZ. Together, these analyses establish a network-level framework for understanding SO–IIS interactions and provide a foundation for future personalized, closed-loop neuromodulatory strategies.

## 2. Materials and Methods

### 2.1 Participants and data acquisition

Six adult patients with drug-resistant focal epilepsy (5 males; mean age: 35 ± 9.5 years) undergoing simultaneous scalp EEG and SEEG monitoring were included in this study. A total of 24 full-day recording days (from a potential 31) were retained after quality screening by excluding recordings with substantial artifacts or without scalp EEG. All participants provided written informed consent, and the study was approved by the University of Chicago Institutional Review Board (protocol 24-1692). Recordings within 5 minutes before to 30 minutes after seizures were excluded from analysis.

Raw sampling rates varied across patients (1024–2048 Hz) and were uniformly downsampled to 256 Hz (MATLAB resample function). All data were originally referenced to FCz. Scalp EEG was acquired using a standard clinical 10–20 montage, with 20 to 26 electrodes across patients. For SO detection in scalp EEG, a common average reference (CAR) was applied to minimize spatial bias, computed as the mean across all scalp channels after excluding mastoid electrodes (M1, M2). After re-referencing, a subset of frontal and prefrontal scalp EEG channels was selected for analysis of PFC activity (F3, F4, F7, F8, F9, F10, Fp1, Fp2, and Fz), following prior research^19^. SEEG depth electrodes were implanted stereotactically to sample the thalamus and the clinically identified SOZ. To minimize volume conduction between adjacent contacts and improve spatial specificity, SEEG signals were re-referenced using a Laplacian scheme: each contact was referenced to the mean of its two immediate physical neighbors on the same electrode shaft. For contacts at the distal or proximal end of a shaft, a bipolar reference (contact minus its single available neighbor) was applied instead. This approach provides the signal-to-noise benefits of bipolar referencing while preserving the anatomical localization of each specific contact^36–38^. Prior to all analyses, 60 Hz and 120 Hz notch filters (4th-order zero-phase Butterworth bandstop filter; applied by MATLAB filtfilt function) was applied to suppress power-line interference.

Thalamic SEEG contacts were localized to multiple nuclei across patients, with predominant coverage of the lateral pulvinar (PuL), centromedian (CM), ventrolateral (VL), anterior (ANT), central lateral (CL), mediodorsal (MD), medial pulvinar (PuM), ventroposterolateral (VPL), ventral anterior (VA), parafascicular (PF), and anterior pulvinar (PuA) nuclei. The SOZ was defined according to clinical documentation, and included the mesial temporal lobe in all participants, as well as temporal neocortical or extratemporal regions in 3 participants. Electrode localization was confirmed by co-registration of post-implantation CT with pre-implantation MRI, normalized to Montreal Neurological Institute space, in BrainStorm^39^. SOZ contacts were assigned to one of three anatomical categories based on their contact-level localization in the AAL3 atlas: mesial temporal, temporal neocortical, or extratemporal. Contact counts after quality screening were 210 PFC (scalp EEG), 177 thalamic (SEEG), and 472 SOZ (SEEG) contacts across the 24 recording days; the 472 SOZ contacts comprised 159 mesial temporal, 253 temporal neocortical, and 60 extratemporal contacts.

### 2.2 Sleep scoring

Sleep stages were scored manually in 30-second epochs by a board certified clinical sleep physician following the American Academy of Sleep Medicine (AASM) guidelines^40^, via scalp EEG signals. Four vigilance states were distinguished: wakefulness (Wake), sleep stage 2 (N2), sleep stage 3 (N3), and combined N2 and N3 (designated NREM in this study). Rapid eye movement (REM) sleep and stage N1 were excluded from all analyses, following prior research^41,42^.

Because scalp EEG does not reliably generate canonical large-amplitude slow oscillations during wakefulness, PFC analyses were restricted to stages N2, N3, and NREM sleep. For thalamic and SOZ contacts, SOs were detected across both wakefulness and sleep (all four vigilance states) using identical morphological criteria; wake-state detections represent slow-wave-like waveforms arising outside the NREM thalamocortical synchronization context^28^, and are referred to as SOs throughout this study for terminological consistency.

### 2.3 SEEG channel polarity determination

Unlike scalp EEG — where the orientation of the cortical current dipole relative to the recording electrode is anatomically constrained and the SO down-state (neuronal silence) invariably manifests as a negative deflection — depth electrodes used for SEEG may record the slow oscillatory dipole from any angle, so that the physiological down-state may appear as either a positive or a negative deflection depending on local electrode geometry^22,43^. Failure to account for this leads to systematic phase errors in SO detection and erroneous assignment of trough time, and artificially inverts apparent SO–IIS coupling relationships in channels where polarity is reversed.

To address this, channel polarity was determined empirically from NREM sleep SO candidates using a gamma-power criterion, following prior research^43^. Genuine NREM slow oscillations have a canonical gamma-band signature: the down-state (neuronal silence) is associated with suppressed gamma power (30–80 Hz), whereas the subsequent up-state is associated with elevated gamma power. For each SEEG channel, the top 25% of sleep SO candidates ranked by absolute trough amplitude were selected. For each of these events, mean gamma power (30–80 Hz, computed via the Hilbert transform after 4th-order Butterworth bandpass filtering) was extracted separately for the negative half-wave (the interval from SO onset to the mid zero-crossing) and the positive half-wave (the interval from the mid zero-crossing to SO end). A channel’s down-state was assigned to the half-wave with lower mean gamma power: if the negative half-wave showed lower gamma power, the channel was retained as-is (down-state = negative deflection, canonical polarity); if the positive half-wave showed lower gamma power, the channel was designated as inverted and all trough/peak assignments were swapped so that the trough always corresponds to the physiological down-state. Crucially, polarity was always estimated from sleep SO candidates only; the same flip decision was then applied to the corresponding wake SO candidates to ensure consistency across vigilance states.

### 2.4 Slow oscillation detection

SOs were detected independently for each anatomical site (PFC, thalamus, and SOZ) using a standard zero-crossing algorithm based on established detection protocols^15,42,44,45^. Raw signals were first bandpass-filtered between 0.1 and 4 Hz using a 4th-order zero-phase Butterworth filter, and candidate SOs were defined as segments between successive positive-to-negative and negative-to-positive zero-crossings. To be classified as an SO, candidates were required to have a negative half-wave duration of 0.25–1.0 s and a total SO duration of 0.5–5.0 s, while excluding candidates that spanned any portion of an artifact or stage N1 or REM sleep. Following candidate detection, an amplitude threshold was applied per channel, retaining only SOs in the top 25% by absolute trough amplitude, computed separately from the respective sleep pool (for sleep SOs) and wake pool (for wake SOs). A kurtosis criterion (waveform kurtosis < 5.0, computed on the bandpass-filtered SO segment) was also applied to exclude events with anomalously non-Gaussian distributions. All analyses were conducted on two parallel SO populations: (1) the AllSO population, comprising all SOs passing the above morphological and epoch criteria; and (2) the gamma-validated population, described in the next section (Section 2.5).

### 2.5 Gamma validation of SO detection fidelity

To mitigate the confounding impact of after-going slow waves following IIS, rather than excluding SOs based on temporal proximity to IIS events, which may obscure physiologically meaningful coupling, we implemented a gamma-based validation step grounded in the physiological definition of slow oscillations. Genuine SOs exhibit a characteristic up-state/down-state structure: neuronal silence during the down-state is associated with reduced gamma-band (30–80 Hz) activity, whereas the subsequent up-state is associated with increased gamma-band activity. This phase-dependent modulation of gamma power has been consistently observed across intracranial and scalp recordings^22,28,30–33^. Accordingly, gamma power was evaluated for candidate SOs to assess whether they preserved canonical up-state/down-state physiology. Gamma power was estimated from the bandpass-filtered (30–80 Hz) signal using the analytic amplitude of the Hilbert transform. Mean gamma power was then extracted separately for the down-state (trough-centered interval) and the subsequent up-state (peak-centered interval). A candidate SO was retained in the gamma-validated population if and only if gamma power during the up-state exceeded that during the down-state, consistent with preserved physiological structure. This approach enables retention of SOs occurring in temporal proximity to IIS (±1 s window) while directly excluding events that do not exhibit canonical slow oscillation dynamics, thereby reducing contamination by IIS-related slow potentials while avoiding reliance on fixed temporal exclusion windows.

### 2.6 IIS detection

IIS were detected exclusively from SOZ SEEG contacts. Each SEEG contact signal was bandpass-filtered between 15 and 50 Hz using a 4th-order zero-phase Butterworth filter. State-specific detection thresholds were computed as six times the standard deviation of the filtered signal within sleep (N2, N3, NREM) and wake states separately, applied only within valid (artifact-free, seizure-free) epochs. Candidate IIS were identified as samples where the absolute filtered amplitude exceeded the threshold. Nearby candidates within 20 samples (∼78 ms at 256 Hz) were grouped, and the peak of the raw (unfiltered) signal within each group was taken as the spike peak (T3). For each candidate T3, the two preceding and two following zero-crossings of the filtered signal were identified (defining landmarks T1, T2–3, T3–4, and T5), and flanking local extrema (T2 and T4) were located. A candidate was retained as a confirmed IIS if: (1) spike duration (T4 − T2) was between 10 and 100 ms; (2) the rise-to-fall asymmetry ratio ((T3 − T2) / (T4 − T3)) was less than 0.75, reflecting the characteristically sharper rising slope of an epileptic spike; and (3) T3 constituted the global maximum of the raw signal amplitude within a ±500 ms window, confirming it as the dominant discharge in the local neighborhood and ensuring that it was not part of an IIS train. The occurrence of a temporally coincident IIS in at least one adjacent contact within 200 ms was required to confirm the candidate event as a true IIS. Visual inspection was performed to confirm detected IIS events. Only IIS occurring in artifact-free, seizure-free epochs were included in SO–IIS coupling analyses.

### 2.7 Peri-event histogram analysis

To characterize the temporal relationship between SOs and IIS, peri-event time histograms of IIS occurrence probability were constructed by aligning events to the SO trough (t = 0), in 50 ms bins spanning a ±1 s window. IIS probability per bin was expressed as a percentage of the total number of SOs contributing to the analysis. A null distribution was derived from stage-matched, event-free control epochs. Statistical significance was assessed using cluster-based permutation testing (1,000 iterations), with surrogate distributions generated by circularly shifting the IIS timestamps relative to SO times, and compared using a cluster-forming threshold of p < 0.05. Results are reported separately for the AllSO and gamma-validated populations, and separately for each origin and vigilance state.

### 2.8 Phase-locking analysis

The instantaneous phase of the SO-band signal was extracted at each IIS timestamp using the Hilbert transform. The SO-band signal was first bandpass-filtered (0.5–1.5 Hz) prior to phase extraction. Phase concentration was quantified using the mean resultant length (MVL), and statistical significance was assessed using the Rayleigh test for circular uniformity^19^. IIS events were included if they occurred within a predefined temporal window around SO troughs (±0.5 s), ensuring physiologically relevant coupling.

### 2.9 SO morphological features and IIS rate correlations

To quantify how SO morphology relates to IIS occurrence at the level of individual recording sites, Spearman rank correlations were computed between five SO morphological features—trough amplitude (μV; negative peak), peak-to-peak amplitude (μV; trough to positive peak), up-slope, down-slope (μV/s; slope of the descending and ascending limbs of the SO waveform), and SO duration (s)—and the IIS rate within a ±1 s window around each SO trough, on a per-SO basis at each contact. Analyses were performed using gamma-validated SOs during N2, N3, and combined

NREM sleep for PFC contacts, and across Wake, N2, N3, and NREM for thalamic and SOZ contacts. Rather than testing significance at individual contacts, the primary objective was to assess the prevalence of morphology–IIS relationships across the recording network.

For each contact, morphological feature, origin, and vigilance state, Spearman correlations were thresholded at p < 0.05, and the proportion of contacts reaching significance was computed. Under the null hypothesis of no association, 5% of contacts are expected to reach this threshold by chance. A one-tailed binomial test was therefore applied to determine whether the observed proportion of significant contacts significantly exceeded this chance level, treating each contact as an independent observation. While contacts are nested within subjects, this analysis is intended to estimate prevalence across spatial sampling rather than to infer subject-level effects. This approach provides a population-level measure of how consistently each SO morphological feature is associated with IIS activity across spatially distributed recording sites. Effect sizes for individual contacts are reported as Spearman ρ, and the dominant direction of association is summarized across the contact population.

### 2.10 Pre-onset phase-amplitude coupling analysis

SOs were classified into two categories based on whether an IIS occurred within the ±1 s peri-trough window: permissive SOs were defined as those associated with at least one IIS within ±1 s of the SO trough; non-permissive SOs were defined as those with no IIS in this window. To characterize the pre-onset network state preceding permissive versus non-permissive SOs, phase–amplitude coupling (PAC) was computed within a pre-onset window defined as the 2 s interval preceding the SO trough (i.e., from −2 s before the SO trough up to SO onset). A 1 s baseline window (−3 to −2 s relative to the SO trough) was used as a spectral reference. PAC was quantified using the mean vector length (MVL) metric, which captures the consistency of phase–amplitude relationships over time^46^. For each cross-frequency pair, the instantaneous phase of the low-frequency component, φ(t), and the amplitude envelope of the high-frequency component, A(t), were extracted using the Hilbert transform after band-pass filtering. These were combined to form a complex-valued signal, as defined in Equation (1):

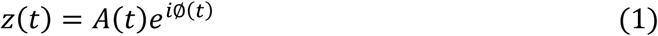

The MVL was then computed as defined in Equation (2):

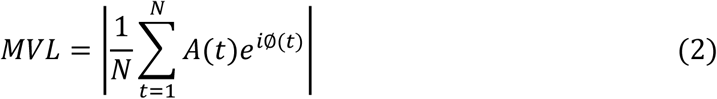

This measure reflects the extent to which high-frequency amplitude is consistently modulated at a specific phase of the low-frequency oscillation, with values near zero indicating weak or random coupling and higher values indicating stronger phase–amplitude coupling. MVL values were baseline-normalized by subtracting the MVL computed in the matched pre-event baseline window.

The following frequency pairs were analyzed: phase bands of δ (0.5–4 Hz), θ (4–8 Hz), and α (8–12 Hz), crossed with amplitude bands of θ (4–8 Hz), α (8–12 Hz), σ (12–16 Hz), β (16–30 Hz), low 𝛤 (𝐿𝛤; 30–59 Hz), and high 𝛤 (𝐻𝛤; 61–119 Hz), yielding 15 cross-frequency feature pairs per contact per SO. All bandpass filters were 4th-order zero-phase Butterworth filters. Origin-level PAC differences between permissive and non-permissive SOs were assessed using paired t-tests at the session level for each frequency pair. t-value matrices comparing PAC between permissive and non-permissive SOs were constructed, with PAC phase frequency on one axis and amplitude frequency on the other, and corrected using the Benjamini–Hochberg false discovery rate (FDR) procedure applied across all frequency pairs within each origin. Significant cells (FDR p < 0.05) indicate frequency pairs where pre-onset PAC systematically differs between permissive and non-permissive SOs.

To characterize the prevalence and heterogeneity of pre-onset PAC differences at the individual structure level (thalamic nuclei and SOZ recording sites), a site-by-feature statistical analysis was conducted. For each recording site, recording day, and PAC frequency pair, PAC values were compared between permissive and non-permissive SOs using a two-sample Welch t-test to account for unequal variances and sample sizes between conditions. To control for multiple comparisons, FDR correction was applied across all tested structure–day–feature combinations within each origin. Effect sizes were quantified using Cohen’s d, and an absolute threshold of |d| ≥ 0.2 was used as a secondary criterion to identify effects of at least small magnitude and exclude trivial differences. Rather than focusing on single-day effects, the primary outcome was defined at the cross-day level to assess robustness. A recording site was classified as predictive if at least 30% of its tested recording days independently exhibited at least one FDR-significant PAC feature difference between permissive and non-permissive SOs. This threshold was chosen to require consistency of the effect across days and reduce the influence of session-specific variability. Mean t-statistics across days were additionally computed for each recording site and PAC feature pair and visualized as heatmaps to summarize the direction and magnitude of effects.

### 2.11 Statistical analysis

All statistical analyses were implemented in MATLAB (MathWorks, version 2025b). Circular statistics (Rayleigh test, MVL, mean resultant direction) were computed using the CircStat toolbox^47^. Multiple comparisons were controlled using the Benjamini–Hochberg FDR procedure throughout. For the peri-event histogram analysis, significance was assessed using nonparametric cluster-based permutation testing (1,000 iterations). Binomial tests were used to determine whether the proportion of thalamic nuclei and SOZ recording sites exceeding a significance threshold was greater than the chance level (5%). Cross-origin comparisons of SO–IIS coupling magnitude were conducted using linear mixed-effects (LME) models with the coupling metric as the dependent variable, brain region (origin) and sleep stage as fixed effects, and subject as a random intercept: Coupling ∼ Origin + Stage + (1|Subject). FDR correction was applied to all post-hoc pairwise comparisons from the LME model. All reported p-values are two-tailed unless otherwise stated. The significance threshold was α = 0.05, and FDR-corrected results were considered significant at p < 0.05.

## 3. Results

### 3.1 Study framework

Figure 1 illustrates the overall experimental framework and data structure used throughout the study. Simultaneous scalp EEG and SEEG recordings enabled concurrent sampling of three anatomically distinct nodes of the thalamocortical–epileptogenic network: the PFC, the thalamus, and the SOZ. All analyses are organized around these three recording origins (Figure 1A). Because reliable large-amplitude slow oscillations are not consistently observed in scalp EEG during wakefulness, PFC analyses were restricted to sleep (N2, N3, and NREM). In contrast, thalamic and SOZ recordings permitted examination of slow oscillatory activity across both wakefulness and sleep (Figure 1B). Figures 1C and 1D provide representative examples of an IIS and an SO from the dataset, serving as a qualitative validation of event detection. Figure 1C illustrates the transient sharp depolarization characteristic of IIS and its propagation across adjacent SEEG contacts. Figure 1E shows the canonical spatial characteristics of scalp-recorded SOs during NREM sleep. This pattern is consistent with well-established canonical properties of slow oscillations during NREM sleep and provides qualitative validation of the SO detection approach^41,42^. Frontal and prefrontal regions exhibit the largest negative trough amplitudes and the highest SO density (events per minute), with a progressive decline toward posterior regions, forming a clear anterior–posterior gradient. For visualization, scalp channels were grouped into five anatomical regions, with the frontal group comprising prefrontal and frontal electrodes. Supplementary Figure S1 shows the same analysis separately for N2 and N3 sleep, demonstrating a consistent spatial pattern across stages, with N3 characterized by a higher overall SO density, particularly over the frontal group, consistent with stronger thalamocortical synchronization during deep sleep.

**Figure 1.**
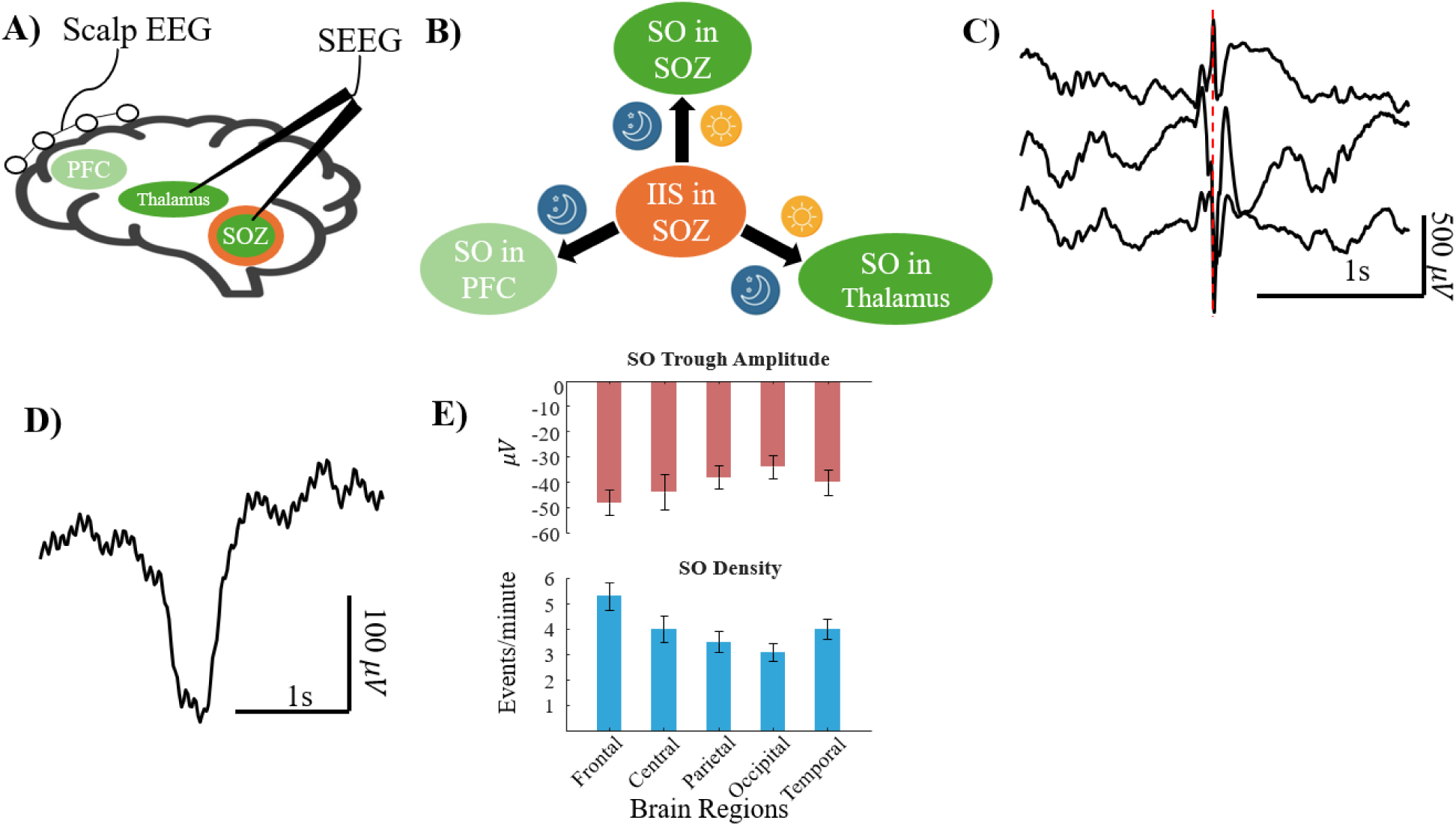
Study framework and experimental overview. **A)** A schematic of the simultaneous recording configuration is presented. Scalp EEG electrodes (prefrontal and frontal montages) capture PFC slow oscillation activity, while depth SEEG electrodes sample thalamic nuclei and the SOZ, enabling concurrent measurement across three anatomically distinct nodes of the thalamocortical–epileptogenic network. **B)** The conceptual framework of the study design is illustrated. SOs are detected independently at each of the three recording origins (PFC, thalamus, SOZ), while IIS are identified exclusively from SOZ contacts. SO–IIS coupling analyses are performed using peri-event histograms and phase-locking analyses across vigilance states (Wake, N2, N3, NREM). PFC analyses are restricted to sleep due to the absence of reliable large-amplitude SOs during wakefulness in scalp recordings. Sleep/wake state icons indicate the states included for each origin. **C)** IIS recorded simultaneously from three adjacent intracranial SEEG contacts in the left parahippocampal region of the SOZ, showing a high-amplitude sharp transient with consistent peak timing across contacts (red dashed line). **D)** Representative SO recorded from a scalp EEG frontal channel (Fz) during stage N3 sleep, illustrating canonical SO morphology characterized by a large-amplitude negative trough followed by a positive peak. **E)** Scalp SO spatial characteristics during NREM sleep (N2+N3), shown as mean (± standard error of the mean [SEM]) SO trough amplitude (μV, upper panel) and SO density (events/min, lower panel) across five scalp regions (Frontal, Central, Parietal, Occipital, Temporal). Both measures show a frontal predominance, with frontal channels exhibiting the most negative trough amplitudes and the highest SO density, and an anteroposterior declining gradient consistent with canonical SO characteristics.

### 3.2 SO–IIS temporal coupling across origins and vigilance states

Peri-event time histograms aligned to the SO trough (50 ms bins, ±1 s window) revealed significant temporal coupling between SOs and IIS across all three recording origins, with coupling strength, and temporal precision varying systematically by origin, and distinct patterns of vigilance-state dependence observed across regions. Phase-locking analyses using the Rayleigh test confirmed origin-level phase concentration in PFC.

#### PFC (NREM)

In the PFC (210 contacts), IIS probability within the peri-SO window was globally elevated relative to the event-free null distribution. A prominent pre-trough IIS peak was observed at t = −0.22 s, reaching 2.6% of IIS probability in the AllSO population. Origin-level phase locking was highly significant (MVL = 0.49, Rayleigh z = 50.37, p < 0.0001), with the preferred IIS phase on the descending limb of the SO (Figure 2A). Following gamma-validation, the pre-trough peak was attenuated to 1.7%, and phase-locking strength was reduced (MVL = 0.34, Rayleigh z = 23.66, p < 0.0001), but the temporal pattern and descending-limb phase preference were fully preserved (Figure 2B). Coupling was detectable in both N2 and N3 sleep (Supplementary Figure S2A).

**Figure 2.**
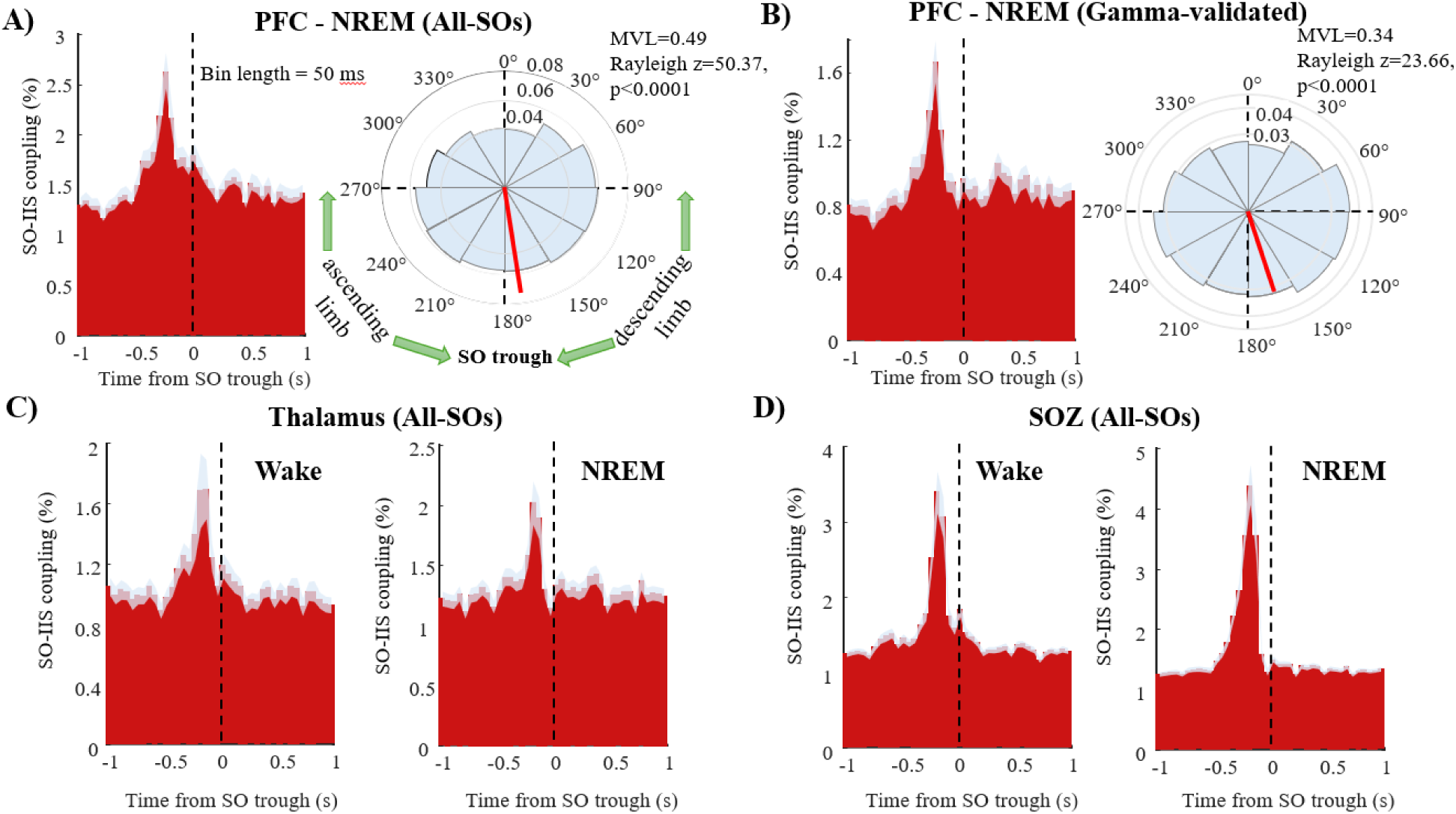
Peri-event SO–IIS coupling histograms and phase analysis across recording origins. **A)** In the PFC during NREM sleep (All SOs; N = 210 contacts), peri-event time histograms (50 ms bins, ±1 s window) show IIS occurrence probability (%) aligned to the SO trough (t = 0, dashed vertical line). Observed IIS probability is shown in red, with variability across contacts indicated by ±SEM (grey shading). Significant clusters were identified using cluster-based permutation testing (1,000 iterations, p < 0.05). A prominent pre-trough IIS peak is observed at t = −0.22 s, reaching 2.6%. Polar histograms of per-contact preferred phases show strong origin-level phase locking (MVL = 0.49, Rayleigh z = 50.37, p < 0.0001), with a preferred phase on the descending limb of the SO. **B)** In the PFC during NREM sleep after gamma-validation (N = 210 contacts), the same analysis shows that exclusion of SOs temporally associated with IIS reduces the pre-trough peak to 1.7%, while preserving the temporal structure and preferred phase. Phase locking remains statistically significant (MVL = 0.34, Rayleigh z = 23.66, p < 0.0001), indicating that the coupling pattern is not driven by IIS-related contamination. **C)** In the thalamus during Wake (left) and NREM sleep (right) (All SOs; N = 177 contacts), peri-event histograms reveal a consistent pre-trough IIS peak at t = −0.17 s in both states, with similar temporal profiles despite differences in vigilance state. **D)** In the SOZ during Wake (left; N = 472 contacts) and NREM sleep (right; N = 430 contacts), peri-event histograms show the strongest temporally coupling across all origins, with a sharp pre-trough IIS peak at t = −0.17 s reaching 3.4% in Wake and 4.4% in NREM. IIS probability is markedly higher during NREM than Wake.

#### Thalamus (Wake and NREM)

In the thalamus (177 contacts), significant SO–IIS coupling was observed during both wakefulness and NREM sleep. In both vigilance states, a pre-trough IIS peak was identified at t = −0.17 s, with the pre-trough timing signature preserved across Wake and NREM (Figure 2C). This state-independent temporal coupling pattern indicates that thalamic SO–IIS phase organization is not contingent on the NREM synchronization context. Gamma validation confirmed that thalamic coupling was not driven by IIS-contaminated SO detection in either state (Supplementary Figure S3A). Coupling was detectable in both N2 and N3 sleep in the thalamus (Supplementary Figure S2B). Despite the robust peri-event temporal coupling, phase analysis revealed no significant origin-level phase locking in the thalamus across vigilance states. The pooled phase distribution did not reach significance in any state (group MVL = 0.05–0.13, Rayleigh p > 0.08), indicating the absence of a consistent preferred phase at the origin level. However, this does not reflect a lack of phase coupling at the nucleus level. This apparent absence of phase locking reflects heterogeneity across thalamic nuclei and is further resolved at the nucleus level (Section 3.3), where structured phase preferences emerge within specific thalamic nuclei.

#### SOZ (Wake and NREM)

The SOZ exhibited the strongest temporal SO–IIS coupling across all three origins. In the AllSO population, a sharp pre-trough IIS peak at t = −0.17 s reached 3.4% during wakefulness (472 contacts) and 4.4% during NREM sleep (430 contacts). The NREM peak exceeded the Wake peak, consistent with sleep-dependent facilitation of epileptiform activity (Figure 2D). Gamma validation confirmed that SOZ coupling was preserved after exclusion of IIS-contaminated SOs in both states (Supplementary Figure S3B). Coupling was detectable in both sleep stages N2 and N3 (Supplementary Figure S2C). Despite this robust peri-event temporal structure, phase analysis revealed no significant origin-level phase locking in the SOZ across vigilance states. The pooled phase distribution did not reach significance in any state (group MVL= 0.01–0.07, Rayleigh p > 0.3), indicating the absence of a consistent preferred phase at the origin level. However, this does not reflect a lack of phase coupling at the individual recording site level. A large majority of SOZ recording sites exhibited significant phase locking (73–86% of recording sites, depending on vigilance state), with relatively high coupling strength (mean MVL up to 0.38 in N3), indicating strong local phase organization of IIS relative to SOs. The absence of a coherent origin-level phase preference therefore reflects heterogeneity of preferred phases across SOZ contacts and regions, consistent with the anatomical and functional diversity of SOZ locations across patients. Peak IIS rates from peri-event histograms and phase-locking metrics are provided in Supplementary Table S1 and Supplementary Table S2, respectively.

### 3.3 Nucleus-level organization of thalamic SO–IIS coupling

Despite robust peri-event temporal coupling, the absence of origin-level phase locking in the thalamus suggests underlying heterogeneity, motivating nucleus-level analysis of SO–IIS coupling. To characterize the spatial distribution of SO–IIS coupling within the thalamus, peri-event histograms and phase-locking measures were examined separately for each of 11 sampled thalamic nuclei across vigilance states, using gamma-validated SOs. Significant coupling (cluster-based permutation, 1,000 iterations, cluster threshold p < 0.05) was detected in 7 of 11 nuclei: PuL, CL, MD, PuM, CM, VPL, and VL. ANT, Pf, PuA, and VA did not reach cluster significance. Significant coupling was most consistent and strongest during N3 sleep, with Wake-state coupling reaching significance in a subset of nuclei (Figure 3A). Phase preferences differed across nuclei and vigilance states, revealing heterogeneous intra-thalamic organization of the SO–IIS coupling signal. In MD during wakefulness (N = 6 contacts; MVL = 0.848, Rayleigh z = 4.32, p = 0.007), IIS preferentially occurred near the SO peak (0°), indicating that MD-synchronized IIS are most probable during the SO up-state (Figure 3B). In contrast, in PuL during NREM sleep (N = 21 contacts; MVL = 0.380, Rayleigh z = 3.03, p = 0.046), the mean preferred phase was approximately 103°, placing IIS on the down-state of the SO prior to the trough (Figure 3C).

**Figure 3.**
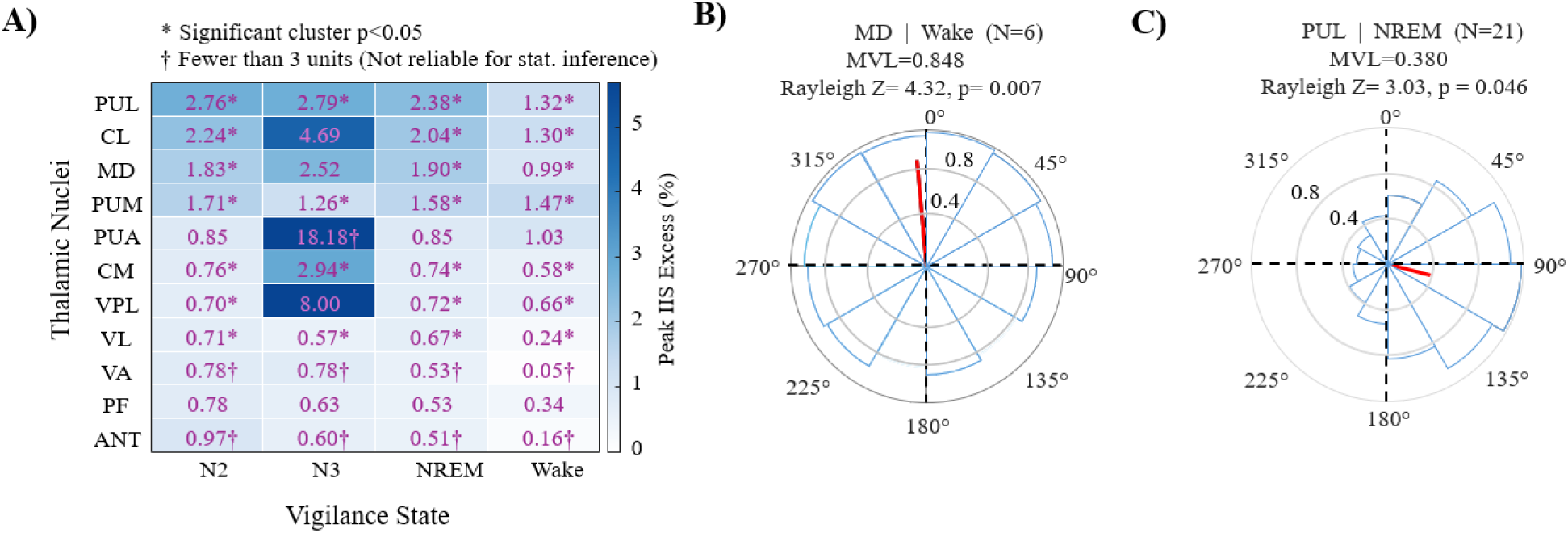
SO–IIS coupling across thalamic nuclei. **A)** Origin-level peri-event SO–IIS coupling across 11 thalamic nuclei was quantified using gamma-validated SOs, with a heatmap showing peak IIS excess (%) across nuclei (rows) and vigilance states (N2, N3, NREM, and Wake). Values represent peak above-baseline IIS probability within the peri-trough window. Asterisks (*) indicate statistically significant clusters identified by cluster-based permutation testing (1,000 iterations, cluster threshold p < 0.05), and daggers (†) denote nuclei with fewer than three contributing contacts, for which statistical inference is unreliable. Significant coupling is observed in 7 of 11 nuclei, with the strongest and most consistent effects during N3 sleep. The lateral pulvinar (PuL), Central Lateral (CL), Mediodorsal (MD), Medial Pulvinar (PuM), Centromedian (CM), Ventroposterolateral (VPL), and Ventral Lateral (VL) nuclei show significant coupling, whereas the anterior (ANT), Parafascicular (Pf), Anterior Pulvinar (PuA), and Ventral Anterior (VA) nuclei do not reach cluster significance; Pf and PuA exhibit non-significant trends. The color scale was capped at the 95th percentile to reduce the influence of outliers on visual contrast. **B)** Phase preference of IIS relative to the SO cycle in the MD during Wake (N = 6 contacts) shows a strong and consistent preferred phase, with polar histograms illustrating the distribution of per-contact preferred phases and the mean resultant vector directed toward the SO peak (0°). This phase organization is reflected in a high coupling strength (MVL = 0.848, Rayleigh z = 4.32, p = 0.007), indicating that IIS in MD preferentially occur near the SO peak (up-state) during wakefulness. **C)** Phase preference of IIS relative to the SO cycle in the PuL during NREM sleep (N = 21 contacts) shows a distinct preferred phase pattern, with the mean resultant vector directed at approximately 103°. Polar histograms illustrate clustering of per-contact preferred phases along the descending phase of the SO, with moderate but significant phase locking (MVL = 0.380, Rayleigh z = 3.03, p = 0.046).

Category-level SO–IIS coupling statistics across the three SOZ anatomical categories are provided in Supplementary Tables S2 and S3. Pre-trough IIS clustering (peak at −175 to −225 ms before the SO trough) was the dominant pattern across all three categories and all vigilance states. Mesial temporal contacts (N = 277) showed the strongest coupling, with peak IIS excess reaching 6.1% during N2, 5.7% during N3, 5.7% during NREM, and 4.0% during wakefulness (all cluster-corrected p < 0.001). Temporal neocortical contacts (N = 117) showed intermediate coupling with peak excess of 3.2% (N2), 2.6% (N3), 2.6% (NREM), and 2.9% (Wake; all p < 0.001). Extratemporal contacts (N = 80) showed the weakest coupling, with peak excess of 2.4% (N2), 1.0% (N3), 2.4% (NREM), and 2.7% (Wake; all p < 0.001). Phase locking across the three categories did not reach significance at the group level (Rayleigh p > 0.5 in all cases), consistent with the heterogeneity of preferred phases across spatially diverse recording sites. Gamma-validated analyses preserved this pattern with modest attenuation (∼10–15% reduction in peak excess), confirming that coupling was not driven by IIS-related slow potentials.

### 3.4 SO morphological features predict IIS rate in the SOZ

To determine whether the morphology of individual PFC SOs modulates IIS occurrence in the SOZ, we computed per-contact Spearman correlations between five SO morphological features — trough amplitude, peak-to-peak amplitude, up-slope, down-slope, and SO duration — and the IIS rate within the ±1 s SO window at each PFC contact, using gamma-validated SOs across N2, N3, and combined NREM sleep. Analyses were performed at the contact level to preserve spatial specificity and capture local variability in SO morphology and IIS coupling. Aggregation at the origin or nucleus level would obscure these relationships by averaging, potentially masking opposing or region-specific effects.

The proportion of PFC contacts showing statistically significant morphology–IIS rate correlations (p < 0.05) significantly exceeded the 5% chance level for trough amplitude, peak-to-peak amplitude, and slope features across all three sleep stages, as assessed by binomial testing in Figure 4. SO duration showed the weakest and least consistent modulation (Figure 4A). The dominant direction of association indicated that steeper up-slopes and larger peak-to-peak amplitudes were associated with increased IIS rate in the SOZ, whereas more negative trough amplitudes and steeper down-slopes were associated with reduced IIS rate (Figure 4B). These associations were replicated in the AllSO population (Supplementary Figure S4), and were also present at thalamic and SOZ contacts (Supplementary Figure S5), though at lower proportions of significant contacts than at PFC.

**Figure 4.**
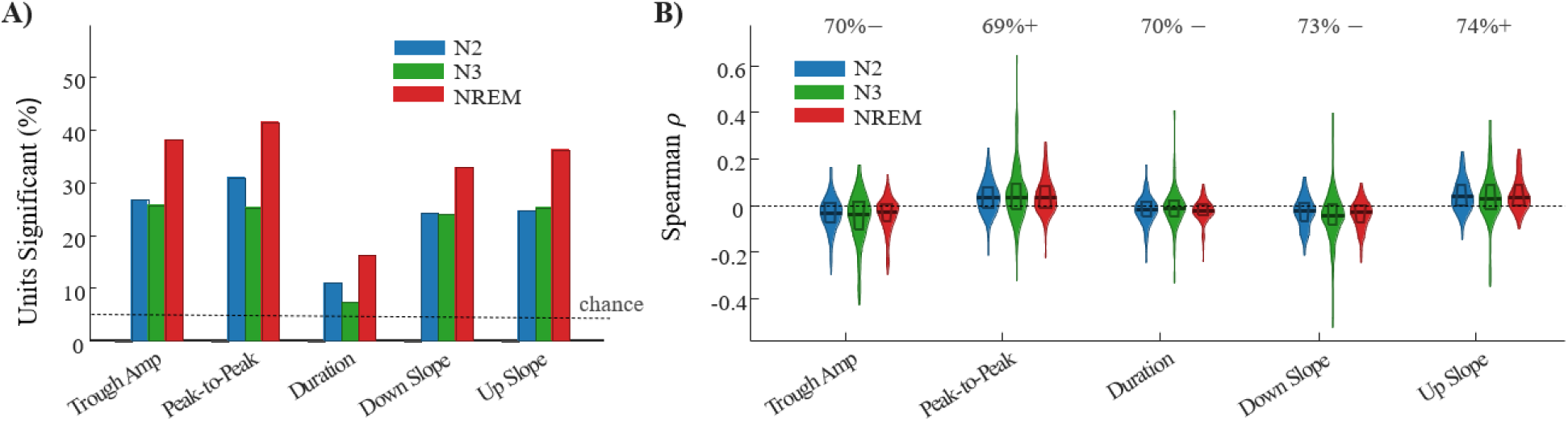
Correlation of SO morphological features in PFC and IIS rate in SOZ during NREM sleep. **A)** The proportion of PFC contacts showing statistically significant Spearman correlations (ρ) between SO morphological features and IIS rate within the ±1 s SO window is shown for N2 (blue), N3 (green), and NREM (red), based on gamma-validated SOs. Significance was assessed at the contact level (p < 0.05), and the dashed horizontal line indicates the 5% chance level. Trough amplitude, peak-to-peak amplitude, and slope features consistently exceed chance across stages, whereas SO duration shows the weakest and least consistent association. **B)** The distribution of per-contact Spearman ρ for each SO morphological feature is shown across PFC contacts, with violin plots illustrating the full distribution and horizontal lines indicating the median and interquartile range. Correlations were computed between SO features measured at each PFC contact and IIS rate in the SOZ within the ±1 s window. The sign of the dominant association is indicated above each feature, where “+” denotes a predominance of positive correlations (increased IIS rate with increasing feature values) and “−” denotes a predominance of negative correlations (decreased IIS rate with increasing feature values).

### 3.5 Pre-onset network state distinguishes permissive from non-permissive SOs

To examine whether permissive SOs are systematically preceded by a distinct network state, we computed PAC within a 2-second pre-onset window preceding each SO trough, using gamma-validated SOs. The analysis timeline, including the baseline and pre-onset windows relative to SO onset and trough, is illustrated in Figure 5A. Paired comparisons of PAC values across frequency bands between permissive and non-permissive SOs (permissive − non-permissive) were computed for PFC, thalamus, and SOZ, all during NREM, with FDR correction applied across all frequency-pair tests within each origin.

**Figure 5.**
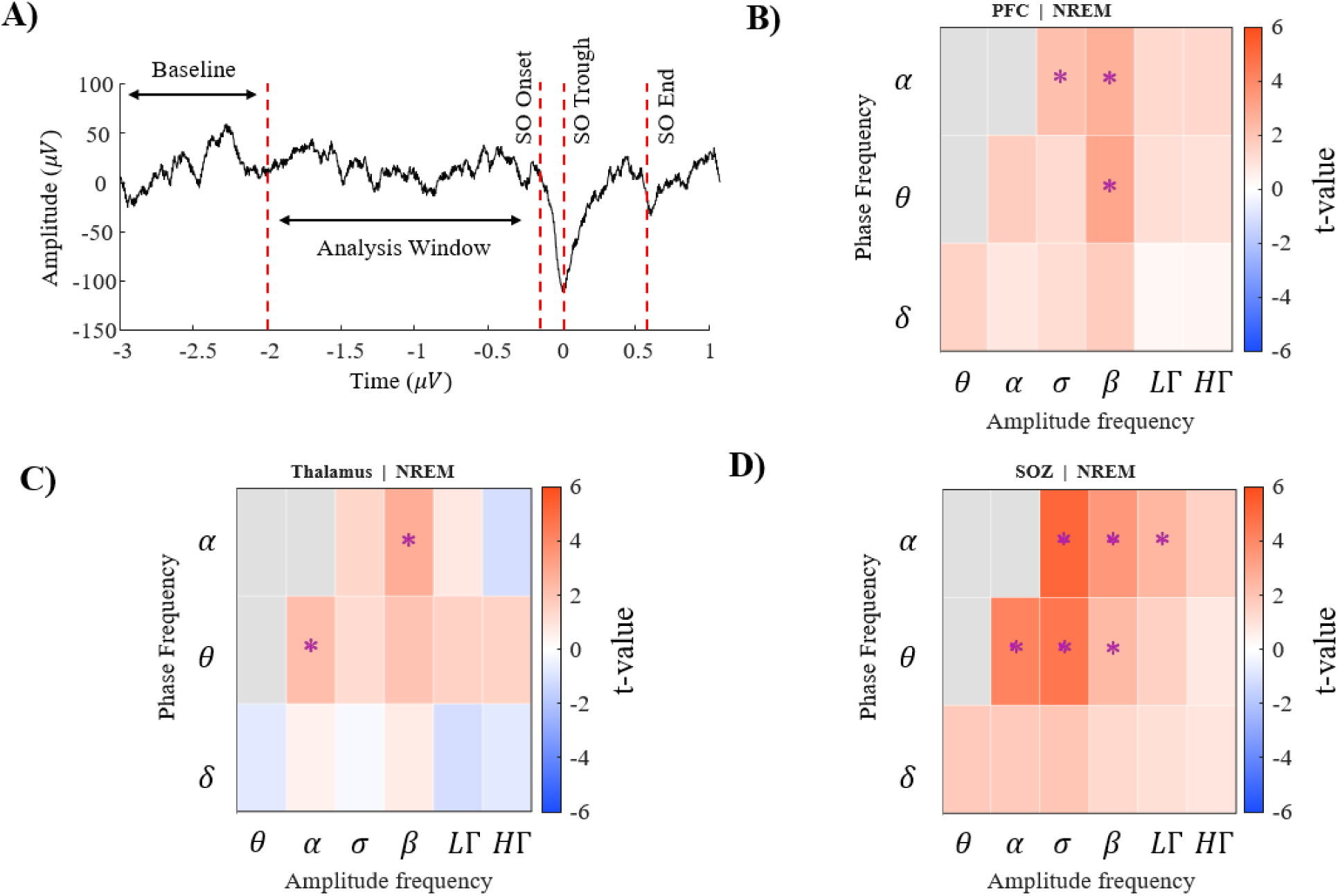
Pre-onset phase-amplitude coupling differs between permissive and non-permissive SOs across recording origins. **A)** The analysis timeline defines the baseline (−3 to −2 s) and pre-onset analysis window (−2 s before trough [t = 0] to SO onset), based on gamma-validated SOs. PAC was computed within the pre-onset window, with the baseline window serving as a spectral reference. SO onset, trough, and end are indicated by red dashed lines. **B)** In the PFC during NREM sleep, the matrix shows paired t-value (permissive − non-permissive SOs) across phase (y-axis) and amplitude (x-axis) frequencies. Significant effects (FDR-corrected, p < 0.05; denoted with asterisks) indicate greater PAC preceding permissive SOs at α-phase modulating β-amplitude, α-phase modulating σ-amplitude, and θ-phase modulating β-amplitude. **C)** In the thalamus during NREM sleep, PAC analysis show significant increases in PAC preceding permissive SOs, primarily at θ-phase modulating α-amplitude and α-phase modulating β-amplitude. **D)** In the SOZ during NREM sleep, PAC analysis reveals the most extensive pre-onset differences, with significant increases in PAC preceding permissive SOs spanning θ/α-phase modulating α/σ/β-amplitude across multiple frequency pairs. This widespread elevation indicates a broadly enhanced pre-onset network state preceding permissive SOs.

#### Origin-level PAC analysis

Significant pre-onset PAC differences were identified in all three origins, with FDR-corrected effects that varied in frequency specificity across the thalamocortical–epileptogenic network. In the PFC, permissive SOs were preceded by significantly greater PAC at α-phase modulating β-amplitude, at α-phase modulating σ-amplitude, and at θ-phase modulating β-amplitude (Figure 5B). This cross-frequency coupling state in the theta–alpha/beta range characterizes the pre-onset period of permissive SOs in the PFC. In the thalamus, significant pre-onset PAC increases were identified at θ-phase modulating α-amplitude and at α-phase modulating β-amplitude, reflecting a partially overlapping but anatomically distinct PAC signature compared to the PFC (Figure 5C). The narrower set of significant frequency pairs in the thalamus is consistent with the more nucleus-specific contributions of thalamic nuclei. The SOZ showed the most extensive PAC differences of the three origins: significant increases in permissive-over-non-permissive pre-onset PAC spanned θ/α-phase modulating α/σ/β-amplitude across multiple frequency pairs surviving FDR correction (Figure 5D). This broad elevation in pre-onset cross-frequency coupling indicates that permissive SOs in the SOZ are systematically preceded by a state of enhanced excitatory network entrainment, detectable up to 2 seconds before SO onset. These results were robustly replicated in the AllSO population.

#### Nucleus- and recording site-level PAC analysis in thalamus and SOZ

PAC analysis at the origin level during wake, using both gamma-validated and AllSO populations, showed no significant effects in either origin (thalamus and SOZ). This motivated us to examine whether SO pre-onset distinguishability in PAC exists during wakefulness at a finer spatial resolution, by dissecting the thalamus and SOZ into their constituent nuclei and recording sites. To characterize which specific thalamic nuclei and SOZ recording sites show robust pre-onset PAC differences at the recording site level, FDR-corrected recording site-level analyses were conducted during wakefulness. A recording site was classified as predictive if ≥30% of recording days showed at least one FDR-significant PAC feature difference between permissive and non-permissive SOs. For the thalamus during wakefulness, Figure 6 displays the nucleus-level mean t-value heatmap showing FDR-corrected PAC differences across 11 thalamic nuclei and 15 PAC feature pairs. PuM emerged as the most predictive thalamic nuclei, meeting the ≥30% recording-day FDR threshold during Wake (40%). CL also met the threshold for Wake. Significant thalamic effects primarily showed permissive > non-permissive directionality (increased PAC preceding permissive events).

**Figure 6.**
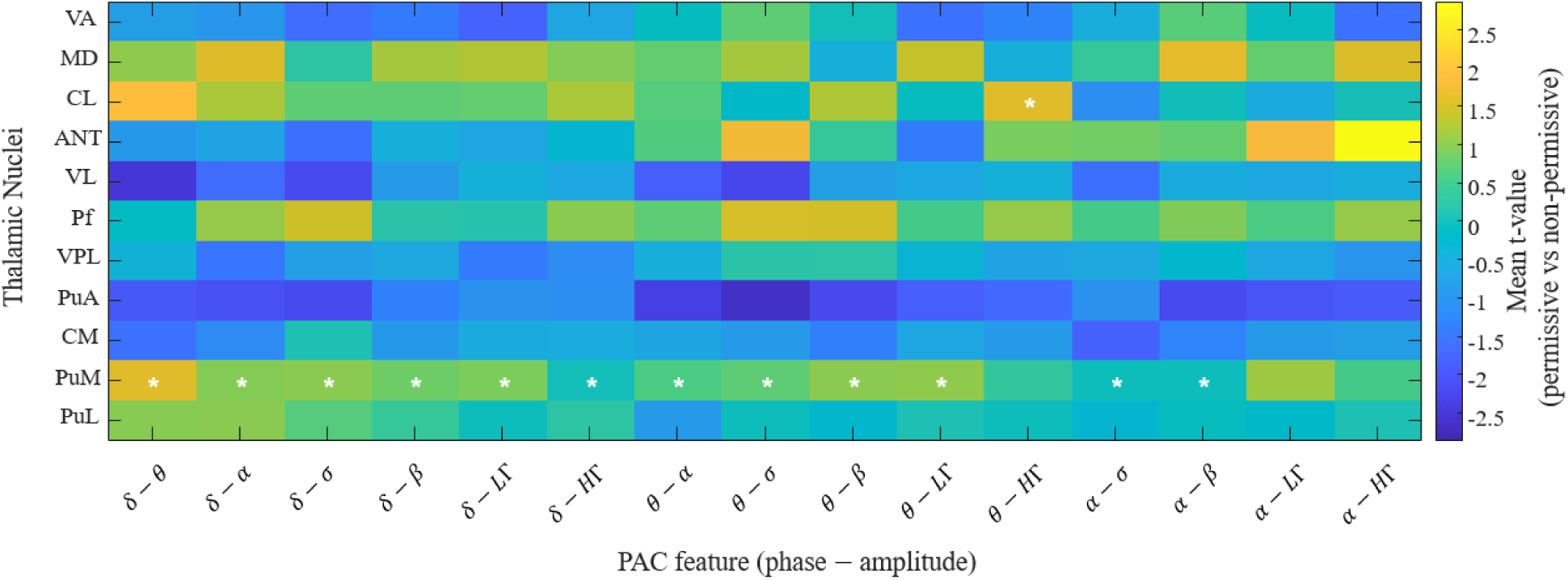
Thalamic nucleus-level PAC t-value heatmap (FDR-corrected) during Wake. Rows represent thalamic nuclei and columns represent PAC features. Color encodes the direction and magnitude of the effect (warm colors: positive t-values, permissive > non-permissive; cool colors: negative t-values, permissive < non-permissive). Asterisks denote nucleus–feature pairs meeting the predictive threshold (FDR p < 0.05 and ≥30% of days). The PuM emerges as the most predictive thalamic nucleus during wakefulness, showing the broadest coverage of significant features. Most significant effects occur in the permissive > non-permissive direction (increased pre-onset PAC preceding permissive SOs).

For the SOZ, category-level FDR-corrected PAC analysis identified predictive pre-onset PAC differences in all three anatomical categories during wakefulness, with the mesial temporal category showing the most extensive and consistent signature. The category-level PAC t-value heatmaps for SOZ during wakefulness are provided in Supplementary Figure S8. During wakefulness, the mesial temporal category met the predictive threshold (≥30% of sessions FDR-significant) across all 15 PAC feature pairs, with uniformly positive directionality (permissive > non-permissive; mean t-statistics ranging from 1 to 2.7), indicating broadly elevated pre-onset cross-frequency coupling preceding IIS-associated SOs. The temporal neocortical category similarly met threshold across all 15 features during wakefulness, with mixed directionality (mean t: −0.9 to 0.8). The extratemporal category met threshold for 6 of 15 features, predominantly showing negative directionality (permissive < non-permissive; mean t: −1.4 to −0.9), suggesting a distinct pre-onset coupling pattern in non-temporal SOZ contacts.

## 5. Discussion

This study provides a comprehensive characterization of SO–IIS coupling across the thalamocortical–epileptogenic network in human focal epilepsy, using simultaneous scalp EEG and SEEG recordings acquired concurrently from three anatomically distinct brain regions across vigilance states. Four principal findings emerge. First, SO–IIS temporal coupling is present across all sampled origins of the thalamocortical–epileptogenic network — the PFC, thalamus, and SOZ — but differs in magnitude, temporal precision, and state-dependence across origins, revealing a hierarchical organization in which the SOZ expresses the strongest and sharpest coupling. Second, coupling at the thalamic level is organized at the resolution of individual nuclei, with distinct phase preferences across nuclei reflecting their heterogeneous connectivity profiles. Third, SO morphological features systematically predict IIS rate in the SOZ, identifying the peak-to-peak amplitude and the slopes of the SO up-state transition as the key morphological determinants of IIS facilitation. Fourth, permissive SOs are preceded by a systematically distinct pre-onset network state characterized by elevated cross-frequency coupling, particularly in the SOZ, that is detectable at a substantial fraction of individual recording sites — providing a distributed biological signature of the IIS-permissive state and a foundational framework for future real-time neuromodulatory approaches.

The finding that SO–IIS coupling magnitude is highest in the SOZ, intermediate in the PFC, and lowest in the thalamus — yet present and statistically significant in all three origins — supports a model in which SO gating of IIS is a network-level phenomenon that is most strongly expressed at the site of IIS generation. The SOZ pre-trough IIS peak, reaching 4.4% during NREM and 3.4% during wakefulness (3.8% and 2.7% after gamma validation, respectively), and markedly exceeding the 1.7% PFC peak following gamma validation, indicates that the SOZ is the primary site of SO–IIS interaction. At the same time, the earlier PFC peak timing (−0.22 s before the SO trough, compared to −0.17 s in both SOZ and thalamus), together with the strong phase locking of IIS to the SO down-state in PFC, suggests that a component of SO–IIS interaction may be modulated, and potentially influenced, by PFC SO activity. In contrast, thalamic coupling (1.1% and 1.3% in wakefulness and NREM, respectively, after gamma validation), despite exhibiting a similar peak IIS timing to the SOZ, may at first glance reflect propagated influences; however, nucleus-level analyses reveal that specific thalamic nuclei also play an active modulatory role in IIS dynamics. The present study is — to our knowledge — the first to compare coupling strength quantitatively across all three origins simultaneously within the same patients and days. The finding that significant SO–IIS coupling is detectable at the scalp level using EEG electrodes has important practical implications, demonstrating that the SO gating signal is not confined to the epileptogenic zone but is accessible via non-invasive recordings.

The SOZ exhibited a clear state-dependent enhancement of coupling, with NREM yielding a markedly higher IIS probability peak (4.4%) than Wake (3.4%), consistent with the well-established sleep-dependent amplification of IIS rates and the heightened synchronization of thalamocortical networks during NREM^21,22^. In contrast, thalamic coupling showed a remarkably similar temporal pattern across Wake and NREM, with the pre-trough IIS peak at t = −0.17 s preserved in both states. These findings suggest that slow oscillatory mechanisms involved in regulating network excitability may not be strictly dependent on the NREM synchronization context, consistent with prior work demonstrating that slow-wave–like activity can occur outside NREM sleep, including during wakefulness, and can modulate epileptic network activity^28^. The N3 sleep stage consistently yielded higher coupling magnitudes and stronger phase locking than N2 across all origins, consistent with the higher amplitude and greater thalamocortical synchrony of N3 SOs^26,48,49^.

The demonstration of significant SO–IIS coupling in 7 of 11 sampled thalamic nuclei — with coupling strength strongest in N3 — provides evidence that the thalamic contribution to the epileptic coupling network is distributed across multiple nuclei. The nuclei showing significant coupling included PuL, PuM, CM, CL, MD, VPL, and VL. This distribution spans nuclei with distinct cortical and subcortical connectivity profiles, including sensory relay, and motor-related thalamic systems. The absence of significant coupling in ANT, Pf, PuA, and VA may reflect either true differences in network involvement or limited sampling within those nuclei. The distinct phase preferences across thalamic nuclei carry mechanistic significance. In MD during wakefulness, IIS preferentially occurred near the SO positive peak, whereas in PuL during NREM sleep, IIS occurred during the descending phase of the SO preceding the trough. These nucleus- and state-specific phase preferences suggest that different thalamic circuits participate in SO–IIS coupling at distinct phases of the oscillatory cycle. Analysis of the SOZ network (Supplementary Tables S2–S3) confirms that pre-trough IIS clustering (−175 to −225 ms before the SO trough) is the dominant pattern across all three SOZ anatomical categories. The strongest coupling was observed within the mesial temporal category (peak NREM excess 5.7%, peak Wake excess 4.0%), consistent with the central role of mesial temporal structures in IIS generation. Temporal neocortical contacts showed intermediate coupling (peak NREM 2.6%) and extratemporal contacts the weakest (peak NREM 2.4%).

The finding that steeper up-slopes and larger peak-to-peak amplitudes of PFC SOs are associated with higher IIS rates in the SOZ is mechanistically interpretable within the up-state excitability framework. A larger peak-to-peak amplitude and a steeper up-slope both reflect a more vigorous and rapid transition from the SO down-state to the up-state, producing a larger and more synchronous burst of activity that, when propagated to the epileptic mesial temporal network, is more likely to recruit IIS generation. Conversely, more negative trough amplitudes and steeper down-slopes were associated with lower IIS rates, indicating a phase-dependent modulation of IIS occurrence across the SO cycle rather than a uniform increase in epileptic activity. The fact that PFC SO morphology (measured at the scalp) predicts IIS rate in the SOZ implies that clinically accessible scalp EEG features of individual SOs could, in principle, serve as a real-time proxy for the likelihood of IIS generation in the SOZ.

The identification of systematic pre-onset PAC differences between permissive and non-permissive SOs across all three recording origins represents a key finding of this study, establishing that the IIS-permissive state is not determined solely by SO morphology or local excitability at the moment of SO occurrence, but is preceded — by at least 2 seconds — by a distinct network state characterized by elevated cross-frequency coupling. The frequency specificity of this pre-onset PAC signature differed across origins in a mechanistically informative manner. In the PFC, the dominant pre-onset signature involved α-phase modulation of β- and σ-amplitude. The SOZ showed the strongest and most extensive pre-onset PAC differences, with significant cross-frequency coupling spanning θ/α-phase modulating α/σ/β-amplitude across multiple frequency pairs that survived FDR correction. Our findings suggest that a comparable multi-frequency interaction characterizes the pre-onset state of permissive SOs in the epileptic network. This pattern may reflect a network configuration associated with an increased likelihood of IIS occurrence, although the causal mechanisms remain to be established. Furthermore, the recording site-level thalamic PAC analysis identified PuM as the most robustly predictive thalamic nucleus during wakefulness. Across nuclei, we observed a predominantly permissive > non-permissive directionality—characterized by increased pre-onset PAC preceding coupling-permissive events. This finding suggests a thalamic synchronization mechanism that may reflect a shift toward increased network excitability associated with periods of elevated IIS susceptibility.

The systematic characterization of pre-onset PAC differences between permissive and non-permissive SOs across the thalamocortical–epileptogenic network provides the empirical foundation and analytic infrastructure for future proactive, closed-loop neuromodulatory approaches. The demonstration that the IIS-permissive state is associated with a distinct, detectable network signature in the SOZ, thalamus, and PFC before SO onset establishes the biological feasibility of a pre-onset detection strategy. A device capable of recognizing this pre-onset PAC signature could, in principle, deliver a targeted neuromodulatory intervention to suppress IIS before they occur. Although established neuromodulatory therapies for epilepsy primarily target seizure reduction rather than IIS suppression per se^50,51^, growing experimental evidence demonstrates that interictal epileptiform activity can be directly modulated, including by closed-loop auditory stimulation during sleep, which has been shown to significantly reduce interictal discharges in human intracranial recordings^52,53^. These findings suggest that IIS-permissive brain states are a viable target for future proactive neuromodulatory strategies. Within the SOZ, mesial temporal contacts, along with delta- and theta-phase coupling to alpha, beta, and gamma amplitude bands, emerge as the most consistent and informative recording sites, providing actionable guidance for the design of such systems. The finding that the mesial temporal and temporal neocortical categories show predominantly permissive > non-permissive directionality (elevated pre-onset PAC preceding IIS-associated SOs), while the extratemporal category shows the inverse pattern, underscores the biological heterogeneity of the epileptic network and the importance of recording site-specific and patient-specific characterization before any intervention is designed. Beyond IIS suppression for cognitive protection, these findings may also be relevant to seizure vulnerability. IIS and seizures are not independent processes, and prior work suggests that their relationship is complex and patient-specific^13^. Identifying SO-gated IIS-permissive states may therefore provide a framework for future studies examining whether such states reflect transient periods of increased epileptogenic network vulnerability and seizure susceptibility^54^.

It is important to note that the pre-onset PAC findings presented here constitute a characterization of the prediction landscape — not a validated real-time classifier. The goal of this analysis is to establish where predictive information is concentrated, and how these features vary across the network and across patients. Translation of these findings into an interventional context requires dedicated future studies in which pre-onset SO state detection is coupled to a therapeutic stimulus — such as transcranial magnetic stimulation, or deep brain stimulation — and evaluated in a controlled clinical design. The present work provides the empirical infrastructure necessary to rationally design and implement such proactive, personalized neuromodulatory strategies for SO-gated IIS suppression in focal epilepsy.

Several limitations of the present study should be acknowledged. First, the sample size is modest, comprising six patients. While this is not atypical for invasive epilepsy research requiring simultaneous scalp EEG and SEEG with coverage of multiple brain regions, it nonetheless limits the generalizability of the findings. Second, there is heterogeneity in SOZ anatomy across patients. Although all patients had mesial temporal SOZ involvement, the specific recording sites and anatomical subregions sampled and the exact seizure network varied across individuals, potentially contributing to variability in coupling magnitude and phase preference. Third, SEEG sampling is inherently sparse and clinically driven, as electrode placement is determined by presurgical requirements rather than systematic research coverage. Accordingly, the absence of significant coupling in certain thalamic nuclei may reflect either true null effects or insufficient sampling. Fourth, the lack of simultaneous invasive PFC recordings represents an additional limitation, as PFC coupling was assessed using scalp EEG, which reflects the spatial average of large cortical territories and has lower spatial resolution and signal-to-noise ratio than intracranial recordings. Finally, the study is observational in design, and no neuromodulatory intervention was performed; therefore, no causal claims can be made regarding the ability of SO modulation to suppress IIS. The pre-onset PAC characterization is intended to map the landscape of predictive information available in the thalamocortical–epileptogenic network — rather than to demonstrate the clinical efficacy of a closed-loop intervention.

This study establishes that SO–IIS coupling in human focal epilepsy is a network-level phenomenon organized across the full thalamocortical–epileptogenic system, with the SOZ expressing the strongest and sharpest temporal coupling, the thalamus contributing a distributed, nucleus-specific, and partially state-independent relay of the coupling signal, and the PFC providing a non-invasive scalp-accessible proxy of the network-wide gating dynamic. SO morphology encodes the strength of coupling at the individual event level, with vigorous up-state transitions facilitating IIS and deep down states suppressing them. Critically, permissive SOs are preceded by a distinct and detectable pre-onset cross-frequency coupling state that is most robustly expressed in the SOZ and is also observable in the PFC, with more limited expression in the thalamus. Together, these findings provide the empirical infrastructure — a network-resolved, recording site-level characterization of SO–IIS coupling and permissive SO pre-onset PAC signatures across invasive and non-invasive modalities — that is necessary to rationally design and implement proactive, personalized neuromodulatory strategies for SO-gated IIS suppression in focal epilepsy.

## Notes

### Competing Interest Statement

The authors have declared no competing interest.

## References

1. Kwan, S.-Y. Sleep in patients with epilepsy. Acta Neurologica Taiwanica 20, 229–231 (2011).

2. Foldvary-Schaefer, N. & Grigg-Damberger, M. Sleep and epilepsy. in vol. 29 419–428 (© Thieme Medical Publishers, 2009).

3. Sunwoo, J.-S. Influence of sleep on seizures and interictal epileptiform discharges in epilepsy. Encephalitis 5, 1 (2024).

4. Moore, J. L., Carvalho, D. Z., St Louis, E. K. & Bazil, C. Sleep and epilepsy: a focused review of pathophysiology, clinical syndromes, co-morbidities, and therapy. Neurotherapeutics 18, 170–180 (2021).

5. Ung, H. et al. Interictal epileptiform activity outside the seizure onset zone impacts cognition. Brain 140, 2157–2168 (2017).

6. Drane, D. L. et al. Interictal epileptiform discharge effects on neuropsychological assessment and epilepsy surgical planning. Epilepsy & Behavior 56, 131–138 (2016).

7. Lambert, I. et al. Hippocampal interictal spikes during sleep impact long-term memory consolidation. Annals of Neurology 87, 976–987 (2020).

8. Okadome, T. et al. The effect of interictal epileptic discharges and following spindles on motor sequence learning in epilepsy patients. Frontiers in Neurology 13, 979333 (2022).

9. Holmes, G. L. Interictal spikes as an EEG biomarker of cognitive impairment. Journal of Clinical Neurophysiology 39, 101–112 (2022).

10. Bernard, C., Frauscher, B., Gelinas, J. & Timofeev, I. Sleep, oscillations, and epilepsy. Epilepsia 64, S3–S12 (2023).

11. Gelinas, J. N. & Khodagholy, D. Interictal network dysfunction and cognitive impairment in epilepsy. Nature Reviews Neuroscience 26, 399–414 (2025).

12. Northcott, E. et al. The neuropsychological and language profile of children with benign rolandic epilepsy. Epilepsia 46, 924–930 (2005).

13. Karoly, P. J. et al. Interictal spikes and epileptic seizures: their relationship and underlying rhythmicity. Brain 139, 1066–1078 (2016).

14. Adamantidis, A. R., Gutierrez Herrera, C. & Gent, T. C. Oscillating circuitries in the sleeping brain. Nature Reviews Neuroscience 20, 746–762 (2019).

15. Massimini, M., Huber, R., Ferrarelli, F., Hill, S. & Tononi, G. The sleep slow oscillation as a traveling wave. Journal of Neuroscience 24, 6862–6870 (2004).

16. Sanchez-Vives, M. V. Origin and dynamics of cortical slow oscillations. Current Opinion in Physiology 15, 217–223 (2020).

17. Staresina, B. P. et al. Hierarchical nesting of slow oscillations, spindles and ripples in the human hippocampus during sleep. Nature neuroscience 18, 1679–1686 (2015).

18. Helfrich, R. F. et al. Bidirectional prefrontal-hippocampal dynamics organize information transfer during sleep in humans. Nature communications 10, 3572 (2019).

19. Schreiner, T., Kaufmann, E., Noachtar, S., Mehrkens, J.-H. & Staudigl, T. The human thalamus orchestrates neocortical oscillations during NREM sleep. Nature communications 13, 5231 (2022).

20. Klinzing, J. G. et al. Spindle activity phase-locked to sleep slow oscillations. Neuroimage 134, 607–616 (2016).

21. Steriade, M., Contreras, D. & Amzica, F. Synchronized sleep oscillations and their paroxysmal developments. Trends in neurosciences 17, 201–207 (1994).

22. Frauscher, B. et al. Facilitation of epileptic activity during sleep is mediated by high amplitude slow waves. Brain 138, 1629–1641 (2015).

23. de Guzman, P. H., Nazer, F. & Dickson, C. T. Short-duration epileptic discharges show a distinct phase preference during ongoing hippocampal slow oscillations. Journal of neurophysiology 104, 2194–2202 (2010).

24. Sheybani, L. et al. Slow oscillations open susceptible time windows for epileptic discharges. Epilepsia 62, 2357–2371 (2021).

25. Ye, H. et al. Widespread slow oscillations support interictal epileptiform discharge networks in focal epilepsy. Neurobiology of Disease 191, 106409 (2024).

26. Neske, G. T. The slow oscillation in cortical and thalamic networks: mechanisms and functions. Frontiers in neural circuits 9, 88 (2016).

27. Gent, T. C., Bandarabadi, M., Herrera, C. G. & Adamantidis, A. R. Thalamic dual control of sleep and wakefulness. Nature neuroscience 21, 974–984 (2018).

28. Sheybani, L. et al. Wake slow waves in focal human epilepsy impact network activity and cognition. Nature Communications 14, 7397 (2023).

29. Kural, M. A. et al. Criteria for defining interictal epileptiform discharges in EEG: A clinical validation study. Neurology 94, e2139–e2147 (2020).

30. Sheroziya, M. & Timofeev, I. Moderate cortical cooling eliminates thalamocortical silent states during slow oscillation. Journal of Neuroscience 35, 13006–13019 (2015).

31. Valderrama, M. et al. Human gamma oscillations during slow wave sleep. PloS one 7, e33477 (2012).

32. Dickson, C. T., Biella, G. & de Curtis, M. Slow periodic events and their transition to gamma oscillations in the entorhinal cortex of the isolated Guinea pig brain. Journal of neurophysiology 90, 39–46 (2003).

33. Compte, A. et al. Spontaneous high-frequency (10–80 Hz) oscillations during up states in the cerebral cortex in vitro. Journal of Neuroscience 28, 13828–13844 (2008).

34. Ngo, H.-V. V., Martinetz, T., Born, J. & Mölle, M. Auditory closed-loop stimulation of the sleep slow oscillation enhances memory. Neuron 78, 545–553 (2013).

35. Esfahani, M. J. et al. Closed-loop auditory stimulation of sleep slow oscillations: Basic principles and best practices. Neuroscience & Biobehavioral Reviews 153, 105379 (2023).

36. Yao, D. et al. Which reference should we use for EEG and ERP practice? Brain topography 32, 530–549 (2019).

37. Parish, G. M., Michelmann, S. & Hanslmayr, S. How Should I Re-reference My Intracranial EEG Data? in Intracranial EEG: A Guide for Cognitive Neuroscientists 451–473 (Springer, 2023).

38. Li, G. et al. Optimal referencing for stereo-electroencephalographic (SEEG) recordings. NeuroImage 183, 327–335 (2018).

39. Tadel, F., Baillet, S., Mosher, J. C., Pantazis, D. & Leahy, R. M. Brainstorm: A user-friendly application for MEG/EEG analysis. Computational intelligence and neuroscience 2011, 879716 (2011).

40. Berry, R. B. et al. AASM scoring manual updates for 2017 (version 2.4). Journal of clinical sleep medicine 13, 665–666 (2017).

41. Alipour, M. et al. The Space–Time Organisation of Sleep Slow Oscillations as Potential Biomarker for Hypersomnolence. Journal of Sleep Research e70059 (2025).

42. Malerba, P., Whitehurst, L. N., Simons, S. B. & Mednick, S. C. Spatio-temporal structure of sleep slow oscillations on the electrode manifold and its relation to spindles. Sleep 42, (2019).

43. Ren, G. et al. Association between interictal high-frequency oscillations and slow wave in refractory focal epilepsy with good surgical outcome. Frontiers in Human Neuroscience 14, 335 (2020).

44. Alipour, M., Seok, S., Mednick, S. C. & Malerba, P. A classification-based generative approach to selective targeting of global slow oscillations during sleep. Frontiers in Human Neuroscience 18, 1342975 (2024).

45. Dang-Vu, T. T. et al. Spontaneous neural activity during human slow wave sleep. Proceedings of the National Academy of Sciences 105, 15160–15165 (2008).

46. Canolty, R. T. et al. High gamma power is phase-locked to theta oscillations in human neocortex. science 313, 1626–1628 (2006).

47. Berens, P. CircStat: a MATLAB toolbox for circular statistics. Journal of statistical software 31, 1–21 (2009).

48. Krishnan, G. P. et al. Cellular and neurochemical basis of sleep stages in the thalamocortical network. Elife 5, e18607 (2016).

49. Pereira, M. et al. Sleep neuroimaging: Review and future directions. Journal of sleep research 34, e14462 (2025).

50. Skarpaas, T. L., Jarosiewicz, B. & Morrell, M. J. Brain-responsive neurostimulation for epilepsy (RNS® System). Epilepsy research 153, 68–70 (2019).

51. Kusyk, D. M., Meinert, J., Stabingas, K. C., Yin, Y. & Whiting, A. C. Systematic review and meta-analysis of responsive neurostimulation in epilepsy. World Neurosurgery 167, e70–e78 (2022).

52. Ngo, H.-V. V., Born, J. & Klinzing, J. G. Protocol for spike-triggered closed-loop auditory stimulation during sleep in patients with epilepsy. STAR protocols 3, (2022).

53. Wong, S. M., et al. Closed-loop modulation of sleep in children undergoing intracranial recordings. Cell Reports Medicine 7, (2026).

54. Lundstrom, B. N., Meisel, C., Van Gompel, J., Stead, M. & Worrell, G. Comparing spiking and slow wave activity from invasive electroencephalography in patients with and without seizures. Clinical Neurophysiology 129, 909–919 (2018).

