## Supplementary Material for "Slow Oscillations Gate Interictal Spikes Across the Human Thalamocortical–Epileptogenic Network"

**Supplementary Tables**

**Supplementary Table S1. Peak IIS Excess and Timing by SOZ Anatomical Category and Vigilance State (AllSO and gamma-validated population).** N = number of contacts contributing to the analysis. Peak excess (%) is the mean observed minus mean control IIS probability at the histogram peak bin. Peak time is the bin center (s) relative to the SO trough. Cluster p-values are from sign-flip cluster-based permutation testing (N = 1,000 iterations). Gamma-validated analyses showed consistent results with ~10–15% attenuation of peak excess.

| SOZ Category | Population | Stage | N | Peak IIS Excess (%) | Peak Bin Center (s) | Cluster p-value at Peak |
|---|---|---|---|---|---|---|
| Mesial Temporal | AllSO | N2 | 277 | 6.066742 | -0.175 | <0.001 |
| Mesial Temporal | AllSO | N3 | 181 | 5.783611 | -0.175 | <0.001 |
| Mesial Temporal | AllSO | NREM | 277 | 5.683531 | -0.175 | <0.001 |
| Mesial Temporal | AllSO | Wake | 277 | 4.006887 | -0.175 | <0.001 |
| Mesial Temporal | gamma-validated | N2 | 277 | 5.512459 | -0.175 | <0.001 |
| Mesial Temporal | gamma-validated | N3 | 181 | 5.428542 | -0.175 | <0.001 |
| Mesial Temporal | gamma-validated | NREM | 277 | 5.12011 | -0.175 | <0.001 |
| Mesial Temporal | gamma-validated | Wake | 277 | 3.377635 | -0.175 | <0.001 |
| Temporal Neocortical | AllSO | N2 | 117 | 3.162721 | -0.225 | <0.001 |
| Temporal Neocortical | AllSO | N3 | 57 | 2.58374 | -0.325 | <0.001 |
| Temporal Neocortical | AllSO | NREM | 117 | 2.646162 | -0.225 | <0.001 |
| Temporal Neocortical | AllSO | Wake | 117 | 2.910697 | -0.175 | <0.001 |
| Temporal Neocortical | gamma-validated | N2 | 117 | 2.572243 | -0.225 | <0.001 |
| Temporal Neocortical | gamma-validated | N3 | 57 | 2.276155 | -0.325 | <0.001 |
| Temporal Neocortical | gamma-validated | NREM | 117 | 2.074311 | -0.225 | <0.001 |
| Temporal Neocortical | gamma-validated | Wake | 117 | 2.04438 | -0.175 | <0.001 |
| Extratemporal | AllSO | N2 | 80 | 2.417926 | -0.225 | <0.001 |
| Extratemporal | AllSO | N3 | 32 | 1.024955 | -0.725 | <0.001 |
| Extratemporal | AllSO | NREM | 80 | 2.377205 | -0.225 | <0.001 |
| Extratemporal | AllSO | Wake | 80 | 2.732885 | -0.225 | <0.001 |
| Extratemporal | gamma-validated | N2 | 80 | 2.040778 | -0.225 | <0.001 |

| | | | | | | |
|---|---|---|---|---|---|---|
| Extratemporal | gamma-validated | N3 | 32 | 0.750845 | -0.725 | <0.001 |
| Extratemporal | gamma-validated | NREM | 80 | 2.009892 | -0.225 | <0.001 |
| Extratemporal | gamma-validated | Wake | 80 | 2.175209 | -0.225 | <0.001 |

**Supplementary Table S2. SO Phase-Locking Statistics by SOZ Anatomical Category and Vigilance State (AllSO and gamma-validated population).** Group MVL and Rayleigh p-value are computed from the distribution of per-contact preferred phases across all contacts in each category. No category reached significance at the group level (all Rayleigh p > 0.5), consistent with heterogeneous phase preferences across the spatially diverse contacts within each category. Mean per-contact MVL reflects local phase locking strength.

| SOZ Category | Population | Stage | N | Rayleigh p-value | Mean Direction (°) | Mean MVL |
|---|---|---|---|---|---|---|
| Mesial Temporal | AllSO | N2 | 277 | 0.867067 | -24.1455 | 0.336021 |
| Mesial Temporal | AllSO | N3 | 231 | 0.898823 | -59.8451 | 0.419546 |
| Mesial Temporal | AllSO | NREM | 277 | 0.785402 | -23.0946 | 0.334342 |
| Mesial Temporal | AllSO | Wake | 277 | 0.924405 | 51.9955 | 0.277306 |
| Mesial Temporal | gamma-validated | N2 | 277 | 0.586687 | -91.2289 | 0.356461 |
| Mesial Temporal | gamma-validated | N3 | 227 | 0.919967 | -88.1839 | 0.441609 |
| Mesial Temporal | gamma-validated | NREM | 277 | 0.588722 | -102.523 | 0.353956 |
| Mesial Temporal | gamma-validated | Wake | 277 | 0.986526 | -63.2215 | 0.284698 |
| Temporal Neocortical | AllSO | N2 | 117 | 0.876654 | -164.343 | 0.195739 |
| Temporal Neocortical | AllSO | N3 | 77 | 0.890927 | -6.1341 | 0.295103 |
| Temporal Neocortical | AllSO | NREM | 117 | 0.99365 | -70.6953 | 0.189706 |
| Temporal Neocortical | AllSO | Wake | 117 | 0.613648 | 25.85385 | 0.202052 |
| Temporal Neocortical | gamma-validated | N2 | 117 | 0.401563 | -164.331 | 0.20046 |
| Temporal Neocortical | gamma-validated | N3 | 74 | 0.954377 | 150.9605 | 0.2691 |
| Temporal Neocortical | gamma-validated | NREM | 117 | 0.534647 | -154.57 | 0.193357 |
| Temporal Neocortical | gamma-validated | Wake | 117 | 0.901616 | 66.47005 | 0.204647 |
| Extratemporal | AllSO | N2 | 80 | 0.799146 | -153.958 | 0.154182 |
| Extratemporal | AllSO | N3 | 53 | 0.839717 | -149.955 | 0.342737 |
| Extratemporal | AllSO | NREM | 80 | 0.524156 | -176.005 | 0.147422 |
| Extratemporal | AllSO | Wake | 80 | 0.987388 | 90.83547 | 0.164799 |
| Extratemporal | gamma-validated | N2 | 80 | 0.999766 | -64.411 | 0.180928 |

| | | | | | | |
|---|---|---|---|---|---|---|
| Extratemporal | gamma-validated | N3 | 46 | 0.992039 | -99.5063 | 0.265554 |
| Extratemporal | gamma-validated | NREM | 80 | 0.890442 | -175.612 | 0.16729 |
| Extratemporal | gamma-validated | Wake | 80 | 0.827734 | 18.96088 | 0.171803 |

### Supplementary Figures

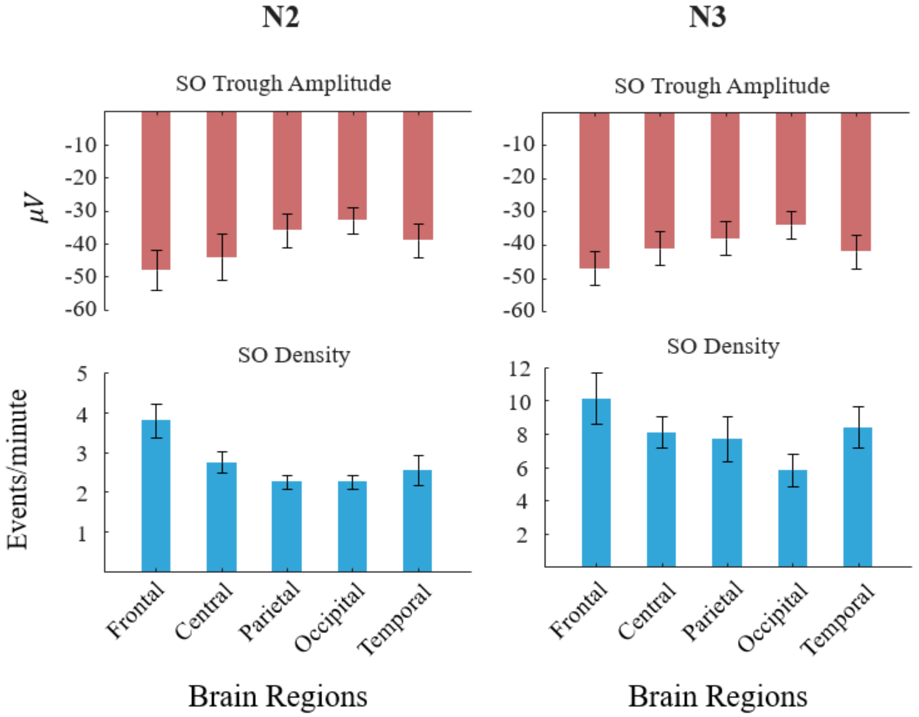


**Supplementary Figure S1. Scalp SO spatial characteristics separately for N2 and N3 sleep.** Mean (±SEM) SO trough amplitude (μV, upper panels) and SO rate (events/min, lower panels) across five scalp electrode regions (Frontal, Central, Parietal, Occipital, Temporal), shown separately for N2 (left column) and N3 (right column). Both sleep stages exhibit a frontal predominance in SO amplitude and density, consistent with the anterior cortical origin of canonical slow oscillations. N3 SOs show markedly higher density than N2 (particularly frontally), consistent with the greater sleep depth of stage N3 facilitating stronger slow oscillatory activity. The anterior–posterior gradient in trough amplitude is present in both stages.

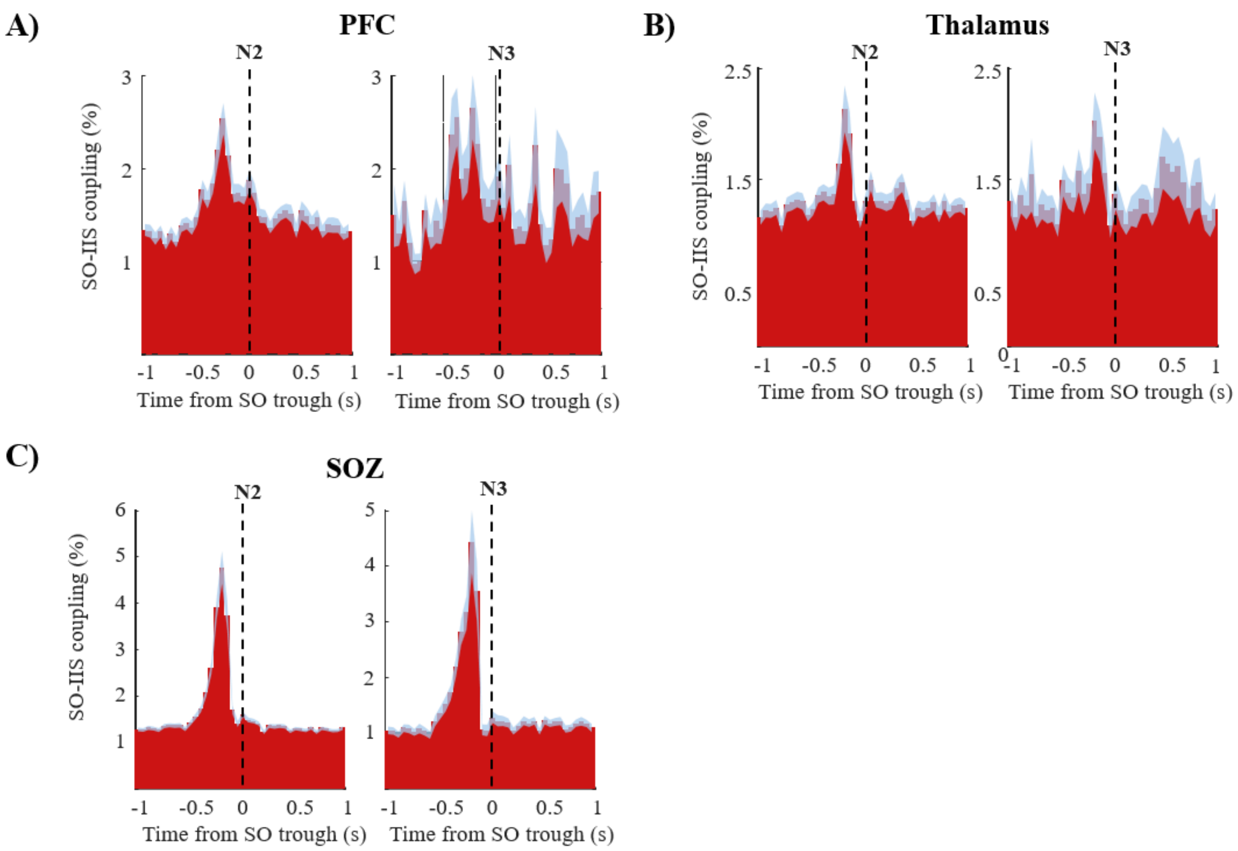


**Supplementary Figure S2. Peri-event SO–IIS coupling histograms for N2 and N3 sleep stages shown separately (AllSO population).** The format and statistical procedures are identical to those in Figure 2. Panels display IIS probability (%, 50 ms bins, ±1 s) aligned to the SO trough ($t = 0$), with shaded envelopes representing ±SEM. Statistically significant clusters identified by cluster-based permutation testing (1,000 iterations, $p < 0.05$) are indicated in red. A) PFC — N2 (left) and N3 (right). B) Thalamus — N2 (left) and N3 (right). C) SOZ — N2 (left) and N3 (right). SO–IIS coupling is evident across all three origins in both N2 and N3, with N3 generally exhibiting larger peak IIS excesses, consistent with enhanced thalamocortical synchrony during deeper sleep and stronger SO-mediated gating of IIS.

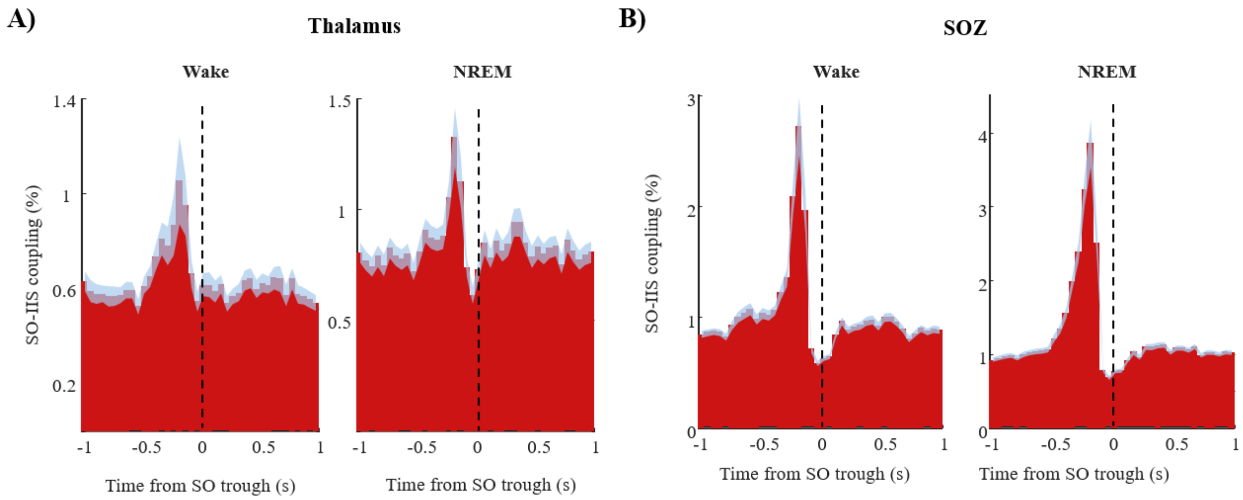


**Supplementary Figure S3. Gamma-validated peri-event SO–IIS coupling histograms for Thalamus and SOZ.** The format and statistical procedures are identical to those in Figure 2. All SOs occurring within ±1.0 s of an IIS were excluded prior to analysis, ensuring that observed coupling is not driven by IIS-related after-going slow waves or contamination of SO detection. **A)** Thalamus — gamma-validated SOs, Wake (left) and NREM (right). The pre-trough IIS peak ($t = -0.17$ s) is preserved after gamma validation in both states, indicating that thalamic SO–IIS coupling reflects genuine phase-locked dynamics rather than an artifact of IIS-contaminated SO detection. B) SOZ — gamma-validated SOs, Wake (left) and NREM (right). The pre-trough coupling peak in the SOZ is similarly preserved following gamma validation, with the NREM peak remaining higher than the Wake peak, consistent with the corresponding AllSO population result in Figure 2D.

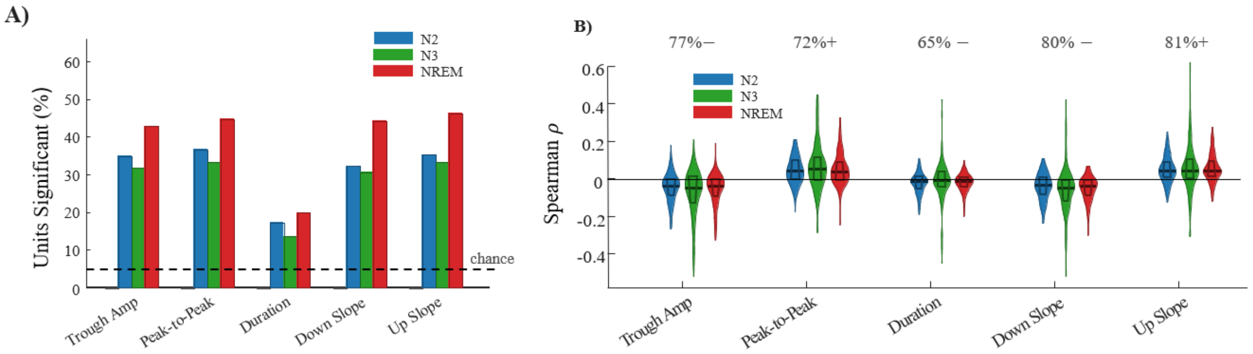


**Supplementary Figure S4. SO morphological predict IIS rate in PFC (AllSO population).** Replication of Figure 4 using all detected SOs. **A)** Percentage of PFC contacts showing significant Spearman correlations ($p < 0.05$) between each SO morphological feature and IIS rate in the SOZ, with the 5% chance level indicated. **B)** Violin plots of per-contact Spearman ρ distributions with annotated effect directions. The overall pattern of associations is consistent with the gamma-validated analysis (Figure 4): trough amplitude, peak-to-peak amplitude, and slope features most reliably predict IIS rate, and the directions of association are preserved. Slightly higher proportions of significant contacts are observed relative to Figure 4, likely reflecting increased statistical power due to the larger number of SO events in the AllSO population. Data are shown for 6 patients across 24 days (PFC, NREM).

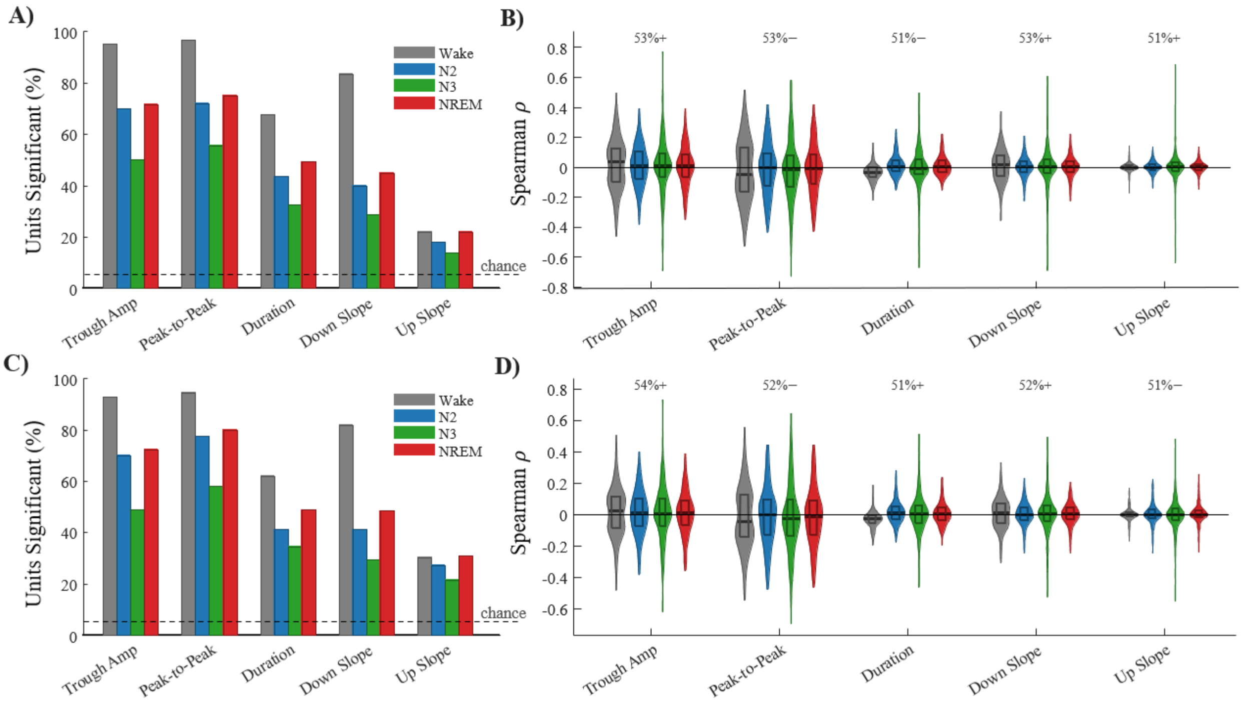


**Supplementary Figure S5. SO morphological predict IIS rate in Thalamus — AllSO and gamma-validated analyses.** Per-contact Spearman correlations (ρ) between SO morphological features measured at individual thalamic contacts and IIS rate in the SOZ within the ±1 s SO window. Panel structure follows Figure 4. **A)** AllSO population — percentage of thalamic contacts with significant Spearman correlations ($p < 0.05$) per SO feature. **B)** AllSO population — violin plots of per-contact ρ distributions with annotated dominant directions. **C)** Gamma-validated population — percentage of significant contacts, same format as A. **D)** Gamma-validated population — violin plots, same format as B. The pattern of thalamic SO morphology associations with IIS rate is broadly consistent across AllSO and gamma-validated populations, though effect magnitudes and the proportion of significant contacts are generally lower than in PFC (Supplementary Figure S4), consistent with the more heterogeneous thalamic contribution to SO–IIS coupling.

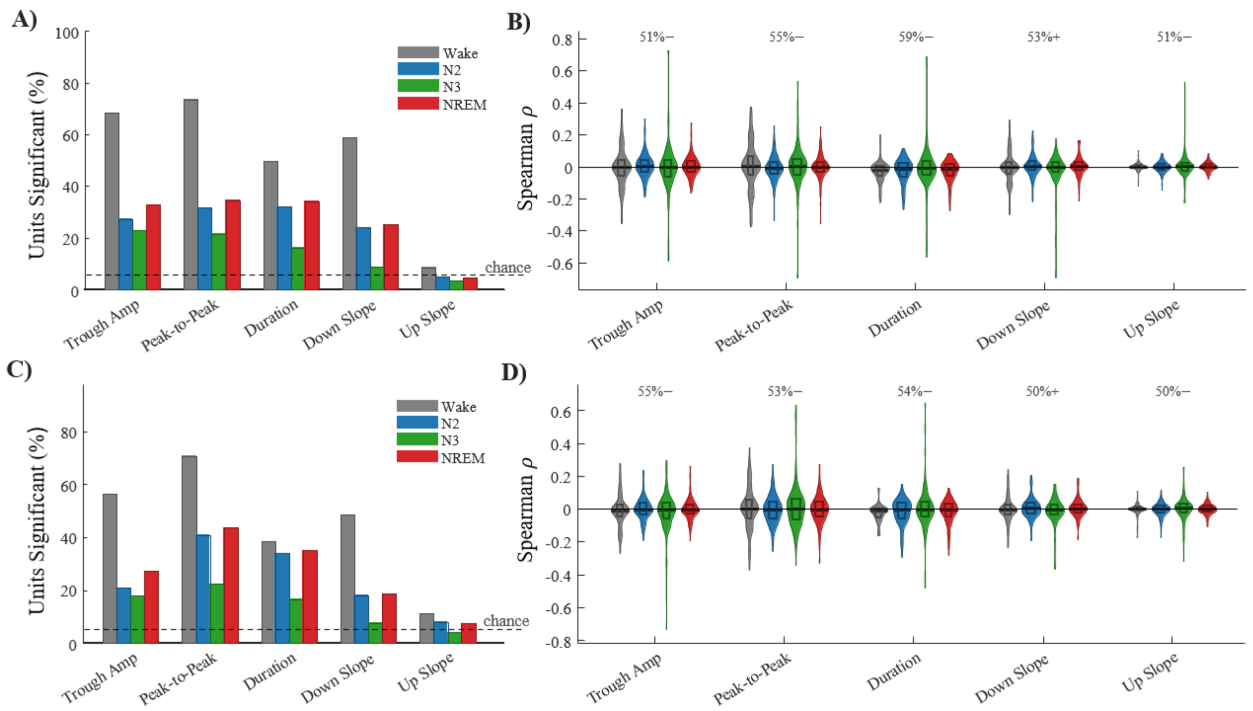


**Supplementary Figure S6. SO morphological predict IIS rate in SOZ — AllSO and gamma-validated analyses.** Per-contact Spearman correlations (ρ) between SO morphological features measured at individual SOZ contacts and IIS rate in the SOZ within the ±1 s SO window. Panel structure follows Figure 4 and Supplementary Figure S5. **A)** AllSO population — percentage of SOZ contacts with significant Spearman correlations ($p < 0.05$) per SO feature. **B)** AllSO population — violin plots of per-contact ρ distributions with annotated dominant directions. **C)** Gamma-validated population — percentage of significant contacts, same format as A. **D)** Gamma-validated population — violin plots, same format as B. SOZ contacts show the highest proportions of significant morphology–IIS correlations across all three origins, with over 50% of contacts reaching significance for the most informative features in the AllSO population. The association pattern is substantially preserved after gamma validation, confirming that the morphological predictors of IIS rate in the SOZ are not driven by IIS-contaminated SOs.

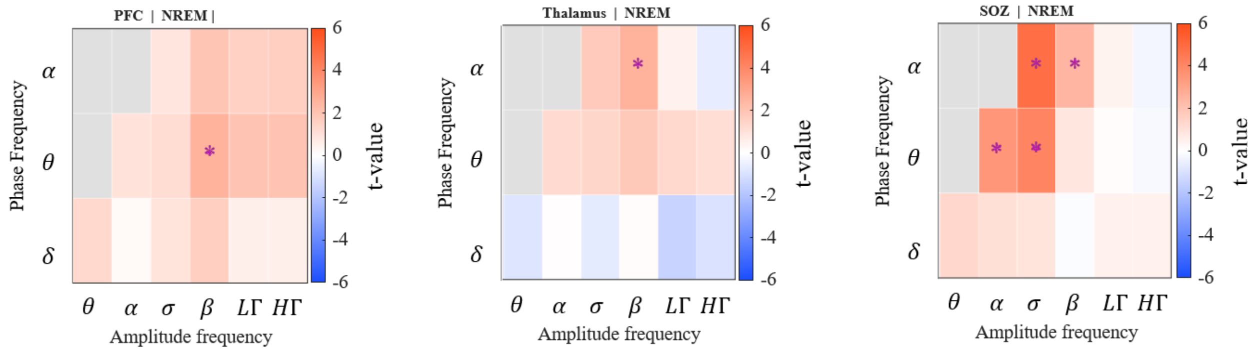


**Supplementary Figure S7. Pre-onset PAC differences between permissive and non-permissive SOs.** Replication of Figure 5 using all detected SOs (AllSO population). PAC matrix show t-values from paired comparisons (permissive − non-permissive), with FDR-corrected significant cells marked (* $p < 0.05$). Red = greater PAC in permissive SOs; blue = greater PAC in non-permissive SOs. **A)** PFC, NREM. **B)** Thalamus, NREM. **C)** SOZ, NREM. The spatial pattern and frequency specificity of PAC differences are broadly consistent with the gamma-validated results (Figure 5), though the number of surviving significant cells is generally higher in the AllSO population due to greater statistical power. The SOZ again shows the most extensive significant PAC differences, spanning θ/α-phase coupling to multiple amplitude bands.

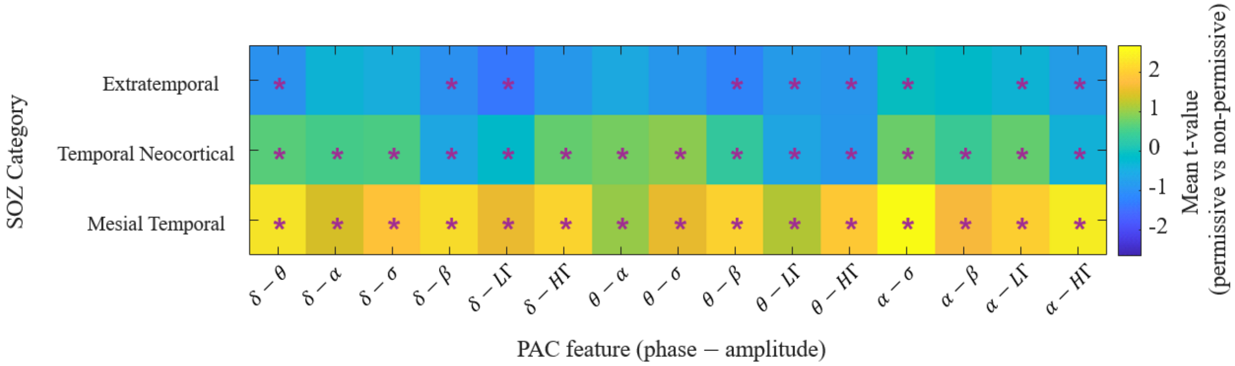


**Supplementary Figure S8.** SOZ category-level PAC t-value heatmap (FDR-corrected) during Wake. Rows represent the three SOZ anatomical categories; columns represent PAC features. Color encodes the mean t-statistic (permissive vs. non-permissive SOs). Asterisks denote category–feature pairs meeting the predictive threshold (FDR $p < 0.05$ in ≥30% of sessions). The mesial temporal category shows the broadest and most consistently positive signature (permissive > non-permissive).
